# When bacteria meet many arms: Autecological insights into *Vibrio pectinicida* FHCF-3 in echinoderms

**DOI:** 10.1101/2025.08.15.670479

**Authors:** Ian Hewson

## Abstract

Sea star wasting (SSW) has been described globally in over 25 species of asteroids. This condition is characterized by body wall lesions, loss of turgor and ray autotomy, which often results in the mortality of specimens. The cause of SSW has remained elusive. A recent report detailing a potential causative agent, *Vibrio pectenicida* FHCF-3 (Prentice et al., 2025), inspired an investigation into its occurrence in available genomic and transcriptomic data from 2013-2015 from wild specimens and those enrolled in experimental incubations. While *Vibrio pectenicida* FHCF-3 16S rRNA gene sequences were detected in abnormal body wall tissues of *Pycnopodia helianthoides* from public aquaria in 2013, they were not detected in grossly normal or abnormal body wall specimens of other species sampled concurrently at sites where mass mortality was observed and from public aquaria. Experimental amendment of *Pisaster ochraceus* with organic matter substrates led to enrichment of *V. pectenicida* FHCF-3 16S rRNAs at the animal-water interface, and that they surged in abundance 24h prior to body wall lesion appearance. However, in this experiment *V. pectenicida* FHCF-3 16S rRNAs were inconsistently detected in coelomic fluid of abnormal specimens, and their abundance at specimen surfaces was inversely related to coelomic fluid detections. Perplexingly, *V. pectenicida* FHCF-3 was detected in abnormal *P. helianthoides* treated with 0.2 µm filtrates of homogenized tissues, but absent in grossly normal heat-treated filtrate controls in prior work. *Vibrio* spp, are copiotrophs that experience rapid growth to dominate microbial communities in plankton and tissues when amended to seawater in a mesocosm experiment. These patterns indicate *V. pectenicida* FHCF-3 might cause abnormalities in *P. helianthoides* under certain conditions, but its growth might be a secondary rather than primary determinant of disease (i.e. it is saprobic or an opportunistic agent). It remains possible that sea star wasting abnormalities in *P. helianthoides* represent a generalized response to bacterial infiltration, driven by a diverse set of bacteria which includes but does not require species such as *V. pectenicida* FHCF-3. Finally, our data suggest that this taxon is not intimately tied to SSW abnormalities in other species. Hence, *V. pectenicida* FHCF-3 may be a driver of a SSW disease in *P. heliathoides*, but cannot be the cause of all SSW across species.

## Introduction

Sea star wasting (SSW) describes a constellation of grossly abnormal signs in asteroids (Asteroidea; Echinodermata) including limb dysplasia, body wall erosions with organ protrusion, loss of turgor, and limb autotomy, which can result in mortality of abnormal specimens (Hewson et al. 2014). SSW has been reported since 1895 (Mead 1898) in over 25 asteroid species asteroids over a wide geographic range (Christensen 1970, Dungan et al. 1982, Eckert et al. 2000, Staehli et al. 2008, Bates et al. 2009, Bucci et al. 2017, Miner et al. 2018, Nunez-Pons et al. 2018, Hewson et al. 2019, Van Volkom et al. 2021, Vergneau-Grosset et al. 2022, Jones & Sewell 2023, Moran et al. 2023, Romer et al. 2025). Beginning in autumn 2013, asteroids along the North American coast experienced mass mortality with SSW signs, beginning on the Pacific Coast of the Olympic peninsula (Washington), followed by the Salish Sea and central California, southern California, Oregon, and Alaska (Hewson et al. 2014). The mass mortality event resulted in the functional extirpation of *Pycnopodia helianthoides* from the Salish Sea (Montecino-Latorre et al. 2016), as well as reduced abundances of other species in subsequent years at routinely monitored sites (Menge et al. 2016, Miner et al. 2018).

The proximal cause of SSW has remained elusive, with several potential infectious and non-infectious etiologies proposed. Early work identified the Sea Star associated Densovirus (SSaDV) as the most likely etiological agent (Hewson et al. 2014). However, subsequent work strongly refuted this hypothesis after re-examination of metagenomic data (Hewson et al. 2018, Hewson et al. 2020a). A non-infectious etiology, the elevated activities of heterotrophic microorganisms at the animal-water interface, has also been advanced (Aquino et al. 2021), but the exact mechanism by which these activities may result in SSW signs in affected stars remains unknown. Temperature, which covaries with several oceanographic parameters, including chlorophyll a and dissolved oxygen, seems to correlate with the elevated occurrence of asteroids with SSW signs in field surveys (Bates et al. 2009, Eisenlord et al. 2016), and cooler fjord waters may represent refugia for some asteroid species during *Pycnopodia helianthoides* mass mortality events (Gehman et al. 2025).

Sea star wasting, at present, has only one case definition amongst the >20 species affected. (Work et al. 2021) described a basal-to-surface process, beginning with inflammation around ossicles, followed by coelomocyte aggregation around these sites before gross signs occur. Histopathology studies of asteroids with SSW signs in 2013-2014 (Hewson et al. 2014a, Bucci et al. 2017) found that lesions were characterized by edema, cleft formation between the outer epidermis and body wall, and necrosis. In no case were cellular inclusions or other signs of invasive infection noted through the use of histochemical stains. Coelomic fluid chemistry of abnormal asteroids is enriched with chloride, protein and coelomocytes (Wahltinez et al. 2020), and the presence of previously undescribed spindle cell coelomocytes (Work et al. 2021). Thorough veterinary hematology revealed bacteria in the coelomic fluid of less than a third of abnormal *Pisaster ochraceus* specimens (Wahltinez et al. 2020).

Recently, Prentice et al. (2025) reported that a SSW condition in *Pycnopodia helianthoides* was associated with metatranscriptomic and V4 16S rRNA amplicon sequences annotated as *Vibrio pectinicida* in libraries prepared from coelomic fluid from wild and tissue homogenate-challenged specimens. Experimental challenge of grossly normal specimens with both tissue homogenates and coelomic fluid from abnormal specimens with SSW signs yielded some abnormalities compatible with SSW (arm twisting and ray autotomy). The authors cultured *V. pectenicida* strain FHCF-3, and challenged grossly normal *P. helianthoides* by injecting this strain into the coelomic cavity of grossly normal asteroids, resulting in limb autotomy and mortality in almost all treated specimens (Prentice et al. 2025). The controls for these experiments comprised both 0.22 µm-filtered coelomic fluid and heat-treated cultures, neither of which resulted in abnormalities, indicating that injection of this bacterial strain may result in limb autotomy in *P. helianthoides* and their demise.

Key questions remain around this finding, especially since the route of transmission in the Prentice et al. (2025) experiment (i.e. injection into coelomic cavity) raises questions about its ability to transmit under more natural settings, i.e. contact exposure. Since the Prentice et al. (2025) study deals exclusively with *P. helianthoides*, it is also unclear whether this result may extend to the other species that are recorded with SSW signs or in other geographic regions (Hewson et al. 2019). Finally, fundamental information about *V. pectenicida*’s mechanism of pathogenesis, its generation of body wall lesions (a key characteristic of SSW), and whether its pathogenesis explains the SSW occurrence in prior years (Hewson et al. 2014, Bucci et al. 2017, Work et al. 2021) remain unknown.

I examined the hypothesis that *V. pectenicida* FHCF-3 is the sole cause of SSW as described in Prentice et al. (2025) by surveying existing databases to evaluate its occurrence in grossly normal and abnormal specimens collected during the asteroid mass mortality of 2013-2014. I also investigated the presence of *V. pectenicida* FHCF-3 16S rRNA in a further suite of asteroid DNA extracts from 2013-2015 using a newly developed PCR assay. I then explored patterns of *V. pectenicida* FHCF-3 16S rRNA amongst 16S rRNA amplicon data from grossly normal and abnormal specimens prepared from experimental incubations. Finally, I examined the response of *Vibrio* spp. to amendment with dead asteroid tissues to constrain their functional roles in echinoderm habitats, especially under decay settings.

## Methods

### 1. Survey of echinoderm samples and sequence databases from earlier mass mortality event

Broad-scale 16S rRNA gene amplicon surveys of asteroids from the 2013 mass mortality were not performed (Hewson et al. 2014, Hewson et al. 2024b). However, virome libraries were prepared from several species of asteroids and holothurians (Hewson et al. 2014, Hewson et al. 2018, Hewson et al. 2020b, Hewson & Sewell 2021), along with transcriptomes of both body wall tissues (Gudenkauf & Hewson 2015) and coelomocytes (Fuess et al. 2015) that are available in the Short-Read Archive (SRA) at NCBI (Table 1).

**Table 1:** Sequence library details that were compared against *Vibrio pectenicida* FHCF-3 copy1.

| Library Accession | Tot Lib Size (reads) | Type of Library | Host | State | Total FHCF-3 copy 1 Matches | Collection Date | Tissue Type | Location | Reference |
| --- | --- | --- | --- | --- | --- | --- | --- | --- | --- |
| SRX8979503 | 2,636,842 | RNA Viral Metagenome | *Apostichopus californius* | Abnormal | 4 | 2016-10-26 | Body Wall | Ketchikan, Alaska | (Hewson et al. 2020b) |
| SRX8979502 | 2,681,239 |  | *Apostichopus californicus* | Grossly Normal | 2 | 2016-10-26 | Body Wall | Ketchikan, Alaska | (Hewson et al. 2020b) |
| SRX8979500 | 2,277,268 |  | *Holothuria scabra* | Grossly Normal | 43 | 2015-12-10 | Body Wall | Dunwich, Australia | (Hewson et al. 2020b) |
| SRX8979498 | 2,275,174 |  | *Holothuria atra* | Grossly Normal | 0 | 2015-12-19 | Body Wall | Heron Island, Australia | (Hewson et al. 2020b) |
| SRX8979505 | 2,741,257 |  | *Holothuria difficilis* | Grossly Normal | 0 | 2015-12-10 | Body Wall | Dunwich, Australia | (Hewson et al. 2020b) |
| SRX8979504 | 2,458,952 |  | *Cucumaria miniata* | Grossly Normal | 0 | 2016-01-07 | Body Wall | Salish Sea | (Hewson et al. 2020b) |
| SRX8979499 | 2,184,535 |  | *Holothuria pardalis* | Grossly Normal | 0 | 2015-12-10 | Body Wall | Dunwich, Australia | (Hewson et al. 2020b) |
| SRX8979497 | 2,246,266 |  | *Stichopus horrens* | Grossly Normal | 0 | 2015-12-10 | Body Wall | Dunwich, Australia | (Hewson et al. 2020b) |
| SRX8979501 | 2,267,391 |  | *Synaptula recta* | Grossly Normal | 0 | 2015-12-10 | Body Wall | Dunwich, Australia | (Hewson et al. 2020b) |
| SRX8825146 | 3,458,388 |  | *Leptasterias sp.* | Grossly Normal | 6 | 2017 | Body Wall | San Franciso, California | (Jackson et al. 2022) |
| SRX876643, SRX876642 | 4127638 |  | *Pycnopodia helianthoides* | Grossly Normal | 4 | 2013-10-26 | Body Wall, Pyloric Caeca, Coelomic Fluid | Seattle Aquarium | (Hewson et al. 2014a) |
| SRX876645, SRX876644 | 4,127,638 |  | *Pycnopodia helianthoides* | Abnormal | 12 | 2013-10-26 | Body Wall, Pyloric Caeca, Coelomic Fluid | Seattle Aquarium | (Hewson et al. 2014a) |
| SRX8825148 | 3,634,059 |  | *Neosmilaster* | Grossly Normal | 0 | 2017 | Body Wall | Palmer Station, Antarctica | (Jackson et al. 2022) |
| SRX8825147 | 845,238 |  | *Mediaster aequalis* | Grossly Normal | 0 | 2016 | Body Wall | Ketchikan, Alaska | (Jackson et al. 2022) |
| SRX3389766 | 2,462,796 |  | *Pisaster ochraceus* | Grossly Normal | 0 | 2013-10-20 | Body Wall | Olympic Peninsula, Washington | (Jackson et al. 2022) |
| SRX3389756,<br>SRX3389754,<br>SRX3389753,<br>SRX3389752,<br>SRX3389751 | 26,309,574 |  | *Pisaster ochraceus* | Grossly Normal | 0 | 2016-02-16 | Body Wall | Salish Sea | (Hewson et al. 2018) |
| SRX8476670,<br>SRX8476668,<br>SRX8476667,<br>SRX8476666,<br>SRX8476665,<br>SRX8476664,<br>SRX8476663,<br>SRX8476662,<br>SRX8476661,<br>SRX8476660,<br>SRX8476659,<br>SRX8476658,<br>SRX8476657,<br>SRX8476656,<br>SRX8476655,<br>SRX8476669, | 26926561 |  | *Pisaster ochraceus* | Abnormal | 498 | 2018-07-01 | Body Wall | Santa Cruz, California | (Hewson et al. 2018) |
| SRX8469595,<br>SRX8469594,<br>SRX8469593 | 4089435 |  | *Stichaster australia,*<br>*Coscinasterias muricata,*<br>*Patiriella sp.* | Grossly Normal | 0 | 2018-01-01 | Body Wall | Auckland, New Zealand | (Hewson & Sewell 2021) |
| SRX3389759 | 4,393,082 | DNA Viral Metagenome | *Acanthaster placi* | Grossly Normal | 15 | 2014-06-06 | Body Wall | Townsville, Australia | (Hewson et al. 2018) |
| SRX3389760 | 2,429,087 |  | *Asterias rubens* | Grossly Normal | 7 | 2014-05-16 | Body Wall | Valparaiso, Chile | (Hewson et al. 2018) |
| SRX3389774 | 5,349,545 |  | *Asterias rubens* | Grossly Normal | 31 | 2014-03-31 | Body Wall | Texel, Netherlands | (Hewson et al. 2018) |
| SRX3389767 | 14,125,699 |  | *Asterias rubens* | Grossly Normal | 8 | 2014-04-19 | Body Wall | Oresund, Netherlands | (Hewson et al. 2018) |
| SRX3389776 | 8,562,699 |  | *Astropecten polyacanthus* | Grossly Normal | 43 | 2014-04-03 | Body Wall | Hong Kong, China | (Hewson et al. 2018) |
| SRX3389775 | 5,183,474 |  | *Echinaster luzonicus* | Grossly Normal | 75 | 2014-03-03 | Body Wall | Okinawa, Japan | (Hewson et al. 2018) |
| SRX875309,<br>SRX875329,<br>SRX875330 | 2818431 |  | *Evasterias troscheli* | Abnormal | 6 | 2013-10-17 | Body Wall, Pyloric Caeca, Coelomic Fluid | Vancouver, British Columbia | (Hewson et al. 2014a) |
| SRX3389771 | 7,176,532 |  | *Helianthus annus* | Grossly Normal | 78 | 2014-05-16 | Body Wall | Valparaiso, Chile | (Hewson et al. 2018) |
| SRX3389758 | 5,298,100 |  | *Henricia ornata* | Grossly Normal | 3 | 2014-05-14 | Body Wall | False Bay, South Africa | (Hewson et al. 2018) |
| SRX8825145 | 3,351,929 |  | *Labidiaster Sp* | Grossly Normal | 4 | 2017 | Body Wall | Palmer Station, Antarctica | (Jackson et al. 2020) |
| SRX3389772 | 5,108,986 |  | *Linckia laevigata* | Grossly Normal | 3 | 2014-05-15 | Body Wall | Chuuk, Micronesia | (Hewson et al. 2018) |
| SRX3389770 | 736,930 |  | *Luidia maculata* | Grossly Normal | 2 | 2014-04-03 | Body Wall | Hong Kong, China | (Hewson et al. 2018) |
| SRX3389755 | 6,025,581 |  | *Marthasterias muizenburg* | Grossly Normal | 8 | 2014-05-14 | Body Wall | False Bay, South Africa | (Hewson et al. 2018) |
| SRX3389769 | 4,657,498 |  | *Marthasterias glacialis* | Grossly Normal | 5 | 2014-05-16 | Body Wall | Valparaiso, Chile | (Hewson et al. 2018) |
| SRX3389768 | 5,465,239 |  | *Ophidiaster sp.* | Grossly Normal | 16 | 2014-03-03 | Body Wall | Okinawa, Japan | (Hewson et al. 2018) |
| SRX3389773 | 4,521,503 |  | *Parvulastera sp.* | Grossly Normal | 19 | 2014-05-14 | Body Wall | False Bay, South Africa | (Hewson et al. 2018) |
| SRX3389757 | 10,552,330 |  | *Patiriella pseudoexigua* | Grossly Normal | 14 | 2014-03-17 | Body Wall | Kaosiung, Taiwan | (Hewson et al. 2018) |
| SRX875314,<br>SRX875319,<br>SRX875320,<br>SRX875325 | 2,805,181 |  | *Pisaster ochraceus* | Grossly Normal | 4 | 2013-10-08 | Body Wall, Pyloric Caeca, Coelomic Fluid | Olympic Peninsula, Washington | (Hewson et al. 2014a) |
| SRX875326,<br>SRX875324,<br>SRX875323,<br>SRX875322,<br>SRX875321 | 2965321 |  | *Pisaster*<br>*ochraceus* | Abnormal | 3 | 2013-10-08 | Body Wall,<br>Pyloric<br>Caeca,<br>Coelomic<br>Fluid | Olympic<br>Peninsula,<br>Washington | (Hewson et al.<br>2014a) |
| SRX3389765,<br>SRX3389764 | 4814415 |  | *Pisaster*<br>*ochraceus* | Grossly<br>Normal | 4 | 2013-10-08 | Body Wall,<br>Pyloric<br>Caeca,<br>Coelomic<br>Fluid | Olympic<br>Peninsula,<br>Washington | (Hewson et al.<br>2014a) |
| SRX875327,<br>SRX875318,<br>SRX875317,<br>SRX875317,<br>SRX875310 | 4,776,177 |  | *Pycnopodia*<br>*helianthoides* | Grossly<br>Normal | 1 | 1905-07-05 | Body Wall,<br>Pyloric<br>Caeca,<br>Coelomic<br>Fluid | Seattle<br>Aquarium | (Hewson et al.<br>2014a) |
| SRX875328,<br>SRX875316,<br>SRX875315,<br>SRX875313,<br>SRX875312 | 4330520 |  | *Pycnopodia*<br>*helianthoides* | Abnormal | 13 | 2013-10-17 | Body Wall,<br>Pyloric<br>Caeca,<br>Coelomic<br>Fluid | Seattle<br>Aquarium | (Hewson et al.<br>2014a) |
| SRX875333 | 523,743 |  | *Patiria miniata* | Abnormal | 0 | 2013-10-17 | Body Wall,<br>Pyloric<br>Caeca,<br>Coelomic<br>Fluid | Monterey<br>Aquarium | (Hewson et al.<br>2014a) |
| SRX875332,<br>SRX875331,<br>SRX875311 | 1270059 |  | *Evasterias*<br>*troscheli* | Grossly<br>Normal | 0 | 2013-10-08 | Body Wall,<br>Pyloric<br>Caeca,<br>Coelomic<br>Fluid | Vancouver,<br>British<br>Columbia | (Hewson et al.<br>2014a) |
| SRX875308,<br>SRX875307 | 2477757 |  | *Solaster*<br>*stimpsoni* | Abnormal | 0 | 2013-10-10 | Body Wall,<br>Pyloric<br>Caeca,<br>Coelomic<br>Fluid | Friday<br>Harbor,<br>Washington | (Hewson et al.<br>2014a) |
| SRX875306 | 503,826 |  | *Dermasterias*<br>*imbricata* | Abnormal | 0 | 2013-10-13 | Body Wall,<br>Pyloric<br>Caeca,<br>Coelomic<br>Fluid | Friday<br>Harbor,<br>Washington | (Hewson et al.<br>2014a) |
| SRX20656813,<br>SRX20656812,<br>SRX20656811,<br>SRX20656810,<br>SRX20656809,<br>SRX20656808,<br>SRX20656807,<br>SRX20656806 | 16,450,843 | Transcriptome | *Pycnopodia*<br>*helianthoides* | Abnormal<br>and GN | 220 | 2020-02-07 | Body Wall,<br>Pyloric<br>Caeca,<br>Coelomic<br>Fluid | Salish Sea | (Schiebelhut et<br>al. 2024) |
| SRX894059,<br>SRX894058,<br>SRX894057 | 122153567 |  | *Pycnopodia*<br>*helianthoides* | Grossly<br>Normal | 71 | 2014 | Coelomocytes | Salish Sea | (Fuess et al.<br>2015) |
| SRX894056,<br>SRX894055,<br>SRX807439,<br>SRX807405 | 44968922 |  | *Pycnopodia*<br>*helianthoides* | Abnormal | 995 | 2013-10-26 | Coelomocytes | Salish Sea | (Fuess et al.<br>2015) |
| SRX906652,<br>SRX906651,<br>SRX906650,<br>SRX612443 | 5,565,373 |  | *Pycnopodia*<br>*helianthoides* | Grossly<br>Normal<br>and<br>Abnormal | >5000 |  | Body Wall | Salish Sea | (Gudenkauf &<br>Hewson 2015) |
| JW523357.1 |  |  | *Tigriopus*<br>*californicus* | Grossly<br>Normal | 2 | 2012 | Whole<br>specimens | Santa Cruz,<br>CA | (Schoville et<br>al. 2012) |

The published *V. pectenicida* FHCF-3 genome bears 8 complete copies of the 16S rRNA gene; 3 copies are 1 - 2nt to the remining 5. I elected to focus on *V. pectenicida* FHCF-3 copy 1 occurrence in this part of the survey, since it had the greatest number of mismatches to relatives in NCBI and would therefore be the best candidate for discriminating this isolate from related species.

Viromes and transcriptomes were queried by BLASTn against the full-length 16S rRNA sequence of *V. pectenicida* FHCF-3 (Copy 1; PQ700178.1). Subject sequences that matched 100% to the query sequence were retrieved, and assembled using the CAP (Contig Assembly Program) (Huang 1992) with minimum overlap of 20 bases and 100% nucleotide identity in overlapping regions. Contigs were then compared against the core_nt database at NCBI by BLASTn (Altschul et al. 1997). Query sequences that were 100% identical to multiple subjects within conserved 16S rRNA gene regions were not considered further since they could not be distinguished from related *Vibrio* spp. Query sequences within variable regions were aligned against close relatives using MUSCLE (Edgar 2004), trimmed for non-overlapping alignment manually, and then subject to phylogenetic analysis using MEGAX (Kumar et al. 2018). We also retrieved sequences matching *V. pectenicida* FHCF-3 copy 1 by BLASTn against the Transcriptome Shotgun Assembly (TSA) and Expressed Sequence Tag (EST) databases at NCBI and included these in our phylogenetic analysis.

In addition to viromes and transcriptomes, we also surveyed 16S rRNA gene amplicon libraries prepared as part of prior work examining the impact of: depleted oxygen on *Asterias forbesi*; organic matter amendment in *Pisaster ochraceus*; and *Pisaster ochraceus* wasting in the absence of external stimuli in aquaria (Aquino et al. 2021) (Table 2). Individual reads matching 100% to the FHCF-3 16S rRNA sequence by BLASTn were retrieved from NCBI. Phylogenetic placement was performed as described above.

**Table 2:** Library information for 16S rRNA amplicons compared against the 16S rRNA of *Vibrio pectenicida* FHCF-3 copy 1.

| Accession Numbers | Treatment | Location | Host | Tissue State | Total Matches | Collection Date | Tissue | Reference |
| --- | --- | --- | --- | --- | --- | --- | --- | --- |
| SRX8657271-SRX8657323 | Depleted O2 through N2 sparging | Ithaca, NY | *Asterias forbesi* | Grossly Normal and Abnormal | 32 | 2019 | Body Wall | (Aquino et al. 2021) |
| SRX8657189-SRX8657415 | Peptone, *Dunaliella salinicola* and coastal POM amendment | Bodega Bay, CA | *Pisaster ochraceus* | Grossly Normal and Abnormal | 10,192 | 2019 | Surface Swab | (Aquino et al. 2021) |
| SRX8657193-SRX8657422 | No external stimuli | Santa Cruz, CA | *Pisaster ochraceus* | Grossly Normal and Abnormal | 2,543 | 2018 | Body Wall | (Aquino et al. 2021) |
| SRX8657189-SRX8657415 | No external stimuli | Santa Cruz, CA | *Pisaster ochraceus* | Abnormal | 53 | 2018 | Body Wall | (Aquino et al. 2021) |
| SRX2753631-SRX2753716 | n/a | Salish Sea | Various Asteroid Species | Grossly Normal | 0 | 2016 | Body Wall | (Jackson et al. 2018) |
| SRX2753631-SRX2753716 | n/a | Moreton Bay and Heron Island, Australia | Various Asteroid Species | Grossly Normal | 0 | 2015 | Body Wall | (Jackson et al. 2018) |
| SRX19779767-SRX19780036 | Depleted O2, glucose, fucose+rhamnose, peptone amendment | Sitka, Alaska | *Apostichopus californicus* | Grossly Normal and Abnormal | 0 | 2021 | Surface Swab | (Crandell et al. 2023) |

### 2. PCR amplification of V. pectenicida in wild specimen tissue extracts from 2013-2016

A PCR primer pair targeting a section of the *V. pectenicida* FHCF-3 16S rRNA gene was designed by first aligning it against all *Vibrio* spp. bacterial 16S rRNAs retrieved from NCBI using MUSCLE (Edgar 2004). *V. pectenicida* and several other species (*V. casei, V. fujianensis, V. natriegens, V. salilacus, V. metschikovii*) 16S rRNA genes contain a 16 nt insertion in the 16S rRNA gene at *E. coli* positions 184 – 205. The insert is heterogeneous between *V. pectinicida* and other *Vibrio* spp. bearing this insert. Hence, the forward primer (Vpec_F) was designed around this region and several nucleotides on the 3’ end (5’-TGTTTGTAATGAACGGGAGCCA-3’).

The reverse primer (Vpec_R) was designed around a variable region at *E. coli* positions 474 – 494 (5’-CTGCAGCTAACGTCAAATGAA-3’). Primer BLAST against NCBI indicated identical matches to *V. pectenicida* FHCF-3 and several other uncultivated *Vibrio* spp. sequences (JQ198184.1, AM990851.1, FJ596526.1, KF941641.1, KF942049.1, JN040578.1, EU253597.1, MK828355.1, HQ690873.1, JF808642.1, JQ199662.1). These represented sequences retrieved from adjacent to marine mammals in aquaria, coastal and offshore plankton, marine sponges, and bivalves. Hence, detections of *V. pectenicida* FHCF-3 via this approach are likely overestimated.

PCR was performed on 91 samples (Table 3) in 15 µl reactions using a BioRad Thermal Cycler. PCR reactions contained 1X One-Taq Quick Load Master Mix, 0.03 µM each of forward and reverse primers and 1 µl of template DNA. Thermal cycling comprised 94°C for 3 minutes, followed by 30 cycles of denaturation at 94°C for 30 s, annealing at 53°C for 30 s and extension at 71°C for 45 s. Cycling was followed by a 5 min finishing step at 71°C, then reactions were held at 15°C until gel electrophoresis. Because specimens were of advanced age (> 10 years), samples were also subject to PCR for the V4 region of the 16S rRNA amplicon employing primers 515F (5’-GTGCCAGCMGCCGCGGTAA-3’) and 806R (5’-GGACTACHVGGGTWTCTAAT-3’) (Caporaso et al. 2010) to validate that they remained viable. The master mix for the 16S rRNA gene was identical in composition to PCR reactions of Vpec_F/Vpec_R. Cycling comprised 5 mins at 95°C, followed by 30 cycles of denaturation at 95°C for 60 s, annealing at 50°C for 60 s and extension at 72°C for 90 s, followed by a finishing step at 72°C for 10 minutes and hold at 12°C until gel electrophoresis.

**Table 3:** Specimen details for PCR amplification of asteroid DNA extracts from 2013-2015. Ampl V4 16S rRNA = Amplicon produced using V4 16S rRNA (prokaryotic-wide) primers; Ampl. Vpec_F/Vpec_R = Amplicon produced using the Vpec_F/Vpec_R primer pair. Note that PCR amplicons using Vpec_F/Vpec_R should be considered overestimates since amplicon sequences of several specimens were from *Vibrio* other than *V. pectenicida* FHCF-3.

| Sample No | Species | Location | Collection Date | Tissue | Tissue State | Ampl. V4 16S rRNA | Ampl. Vpec_F/Vpec_R |
| --- | --- | --- | --- | --- | --- | --- | --- |
| s13 | *Pisaster ochraceus* | Scott Creek, Santa Cruz | 2013-10-08 | Pyloric Caeca | Abnormal |  |  |
| s15 | *Pisaster ochraceus* | Scott Creek, Santa Cruz | 2013-10-08 | Pyloric Caeca | Abnormal | + |  |
| s16 | *Pisaster ochraceus* | Scott Creek, Santa Cruz | 2013-10-08 | Body Wall | Abnormal | + |  |
| s17 | *Pisaster ochraceus* | Scott Creek, Santa Cruz | 2013-10-08 | Pyloric Caeca | Abnormal | + |  |
| s18 | *Pisaster ochraceus* | Scott Creek, Santa Cruz | 2013-10-08 | Body Wall | Abnormal | + | + |
| s19 | *Pisaster ochraceus* | Natural Bridges, Santa Cruz | 2013-09-20 | Pyloric Caeca | Abnormal | + |  |
| s20 | *Pisaster ochraceus* | Natural Bridges, Santa Cruz | 2013-09-20 | Body Wall | Abnormal | + |  |
| s21 | *Pisaster ochraceus* | Natural Bridges, Santa Cruz | 2013-09-20 | Pyloric Caeca | Abnormal | + |  |
| s23 | *Pisaster ochraceus* | Terrace Point, Santa Cruz | 2013-09-20 | Pyloric Caeca | Abnormal |  |  |
| s29 | *Pisaster ochraceus* | Pigeon Point, Santa Cruz | 2013-10-17 | Pyloric Caeca | Abnormal | + |  |
| s31 | *Pycnopodia helianthoides* | Vancouver Aquarium | 2013-10-16 | Body Wall | Abnormal | + | + |
| s43 | *Evasterias troscheli* | Cape Roger Curtis, Vancouver | 2013-10-17 | Body Wall | Abnormal | + |  |
| s44 | *Evasterias troscheli* | Cape Roger Curtis, Vancouver | 2013-10-17 | Pyloric Caeca | Abnormal | + |  |
| s45 | *Evasterias troscheli* | Cape Roger Curtis, Vancouver | 2013-10-17 | Body Wall | Abnormal | + |  |
| s46 | *Evasterias troscheli* | Cape Roger Curtis, Vancouver | 2013-10-17 | Pyloric Caeca | Abnormal | + |  |
| s67 | *Pycnopodia helianthoides* | Friday Harbor, Washington | 2013-10-11 | Pyloric Caeca | Abnormal | + |  |
| s68 | *Evasterias troscheli* | Friday Harbor, Washington | 2013-10-09 | Body Wall | Grossly Normal | + |  |
| s91 | *Pisaster ochraceus* | Olympic National Park | 2013-10-20 | Pyloric Caeca | Abnormal | + |  |
| s92 | *Pisaster ochraceus* | Olympic National Park | 2013-10-20 | Body Wall | Abnormal | + |  |
| s93 | *Pisaster ochraceus* | Olympic National Park | 2013-10-20 | Pyloric Caeca | Abnormal | + |  |
| s95 | *Pisaster ochraceus* | Olympic National Park | 2013-10-20 | Pyloric Caeca | Abnormal |  |  |
| s97 | *Pisaster ochraceus* | Olympic National Park | 2013-09-19 | Body Wall | Grossly Normal | + |  |
| s98 | *Pisaster ochraceus* | Olympic National Park | 2013-09-19 | Pyloric Caeca | Grossly Normal | + |  |
| s110 | *Pisaster ochraceus* | Olympic National Park | 2013-09-19 | Pyloric Caeca | Abnormal | + |  |
| s111 | *Pisaster ochraceus* | Olympic National Park | 2013-09-19 | Body Wall | Abnormal | + |  |
| s112 | *Pisaster ochraceus* | Olympic National Park | 2013-09-19 | Pyloric Caeca | Abnormal | + |  |
| s113 | *Pisaster ochraceus* | Olympic National Park | 2013-09-19 | Body Wall | Abnormal | + |  |
| s114 | *Pisaster ochraceus* | Olympic National Park | 2013-09-19 | Pyloric Caeca | Abnormal | + |  |
| s115 | *Pisaster ochraceus* | Olympic National Park | 2013-09-19 | Body Wall | Abnormal | + |  |
| s116 | *Pisaster ochraceus* | Olympic National Park | 2013-09-19 | Pyloric Caeca | Abnormal | + |  |
| s117 | *Pisaster ochraceus* | Olympic National Park | 2013-09-19 | Body Wall | Abnormal | + |  |
| s118 | *Pisaster ochraceus* | Olympic National Park | 2013-09-19 | Pyloric Caeca | Abnormal | + |  |
| s130 | *Evasterias troscheli* | Cates Reef Park, Vancouver | 2013-10-29 | Body Wall | Abnormal |  |  |
| s131 | *Evasterias troscheli* | Cates Reef Park, Vancouver | 2013-10-29 | Pyloric Caeca | Abnormal | + |  |
| s144 | *Pycnopodia helianthoides* | Seattle Aquarium | 2013-10-26 | Pyloric Caeca | Abnormal | + | + |
| s146 | *Pycnopodia helianthoides* | Seattle Aquarium | 2013-10-26 | Pyloric Caeca | Abnormal | + | + |
| s147 | *Pycnopodia helianthoides* | Seattle Aquarium | 2013-10-26 | Body Wall | Abnormal | + | + |
| s148 | *Pycnopodia helianthoides* | Seattle Aquarium | 2013-10-26 | Pyloric Caeca | Abnormal | + | + |
| s149 | *Pycnopodia helianthoides* | Seattle Aquarium | 2013-10-26 | Body Wall | Abnormal | + | + |
| s150 | *Pycnopodia helianthoides* | Seattle Aquarium | 2013-10-26 | Pyloric Caeca | Abnormal | + | + |
| s151 | *Pycnopodia helianthoides* | Seattle Aquarium | 2013-10-26 | Body Wall | Abnormal | + | + |
| s152 | *Pycnopodia helianthoides* | Seattle Aquarium | 2013-10-26 | Pyloric Caeca | Abnormal | + | + |
| s153 | *Evasterias troscheli* | Seattle Aquarium | 2013-10-26 | Body Wall | Abnormal | + |  |
| s154 | *Evasterias troscheli* | Seattle Aquarium | 2013-10-26 | Pyloric Caeca | Abnormal | + | + |
| s155 | *Pycnopodia helianthoides* | Seattle Aquarium | 2013-11-01 | Body Wall | Abnormal | + |  |
| s156 | *Pycnopodia helianthoides* | Seattle Aquarium | 2013-11-01 | Pyloric Caeca | Abnormal | + |  |
| s157 | *Pycnopodia helianthoides* | Seattle Aquarium | 2013-11-01 | Body Wall | Abnormal | + | + |
| s158 | *Pycnopodia helianthoides* | Seattle Aquarium | 2013-11-01 | Pyloric Caeca | Abnormal | + | + |
| s159 | *Pycnopodia helianthoides* | Seattle Aquarium | 2013-11-01 | Body Wall | Abnormal | + | + |
| s160 | *Pycnopodia helianthoides* | Seattle Aquarium | 2013-11-01 | Pyloric Caeca | Abnormal | + | + |
| s163 | *Pycnopodia helianthoides* | Seattle Aquarium | 2013-11-01 | Body Wall | Abnormal | + | + |
| s165 | *Pycnopodia helianthoides* | Vancouver Aquarium | 2013-11-01 | Body Wall | Abnormal | + | + |
| s167 | *Pycnopodia helianthoides* | Vancouver Aquarium | 2013-11-01 | Body Wall | Abnormal | + | + |
| s169 | *Pycnopodia helianthoides* | Vancouver Aquarium | 2013-11-01 | Body Wall | Abnormal | + | + |
| s171 | *Pycnopodia helianthoides* | Vancouver Aquarium | 2013-11-17 | Body Wall | Abnormal | + | + |
| s318 | *Pycnopodia helianthoides* | Vancouver Aquarium | 2013-12-31 | Body Wall | Abnormal |  |  |
| s327 | *Pycnopodia helianthoides* | Vancouver Aquarium | 2013-11-01 | Body Wall | Abnormal |  |  |
| s328 | *Pycnopodia helianthoides* | Vancouver Aquarium | 2013-11-17 | Body Wall | Abnormal |  |  |
| s334 | *Pycnopodia helianthoides* | Vancouver Aquarium | 2013-11-01 | Body Wall | Abnormal |  |  |
| s394 | *Pycnopodia helianthoides* | Salisbury, Washington | 2014-01-29 | Body Wall | Abnormal |  |  |
| s581 | *Pycnopodia helianthoides* | Redondo Bay, Washington | 2013-11-19 | Body Wall | Abnormal |  |  |
| s584 | *Pycnopodia helianthoides* | Redondo Bay, Washington | 2013-11-19 | Body Wall | Abnormal |  |  |
| s595 | *Pycnopodia helianthoides* | Carpentaria, California | 2014-02-25 | Body Wall | Abnormal |  |  |
| s596 | *Pycnopodia helianthoides* | Redondo Bay, Washington | 2014-01-20 | Body Wall | Abnormal | + | + |
| A22 | *Pycnopodia helianthoides* | Dutch Harbor, Alaska | 2015-03-11 | Tube Foot | Grossly Normal | + |  |
| A24 | *Pycnopodia helianthoides* | Dutch Harbor, Alaska | 2015-03-11 | Tube Foot | Grossly Normal | + |  |
| A25 | *Pycnopodia helianthoides* | Dutch Harbor, Alaska | 2015-03-11 | Tube Foot | Grossly Normal | + |  |
| A26 | *Pycnopodia helianthoides* | Dutch Harbor, Alaska | 2015-03-11 | Tube Foot | Grossly Normal | + |  |
| A38 | *Pycnopodia helianthoides* | Dutch Harbor, Alaska | 2015-03-11 | Tube Foot | Grossly Normal | + |  |
| A39 | *Pycnopodia helianthoides* | Dutch Harbor, Alaska | 2015-03-11 | Tube Foot | Grossly Normal | + |  |
| A40 | *Pycnopodia helianthoides* | Dutch Harbor, Alaska | 2015-03-11 | Tube Foot | Grossly Normal |  |  |
| A41 | *Pycnopodia helianthoides* | Dutch Harbor, Alaska | 2015-03-11 | Tube Foot | Grossly Normal |  |  |
| A42 | *Pycnopodia helianthoides* | Dutch Harbor, Alaska | 2015-03-11 | Tube Foot | Grossly Normal | + |  |
| A43 | *Pycnopodia helianthoides* | Dutch Harbor, Alaska | 2015-03-11 | Tube Foot | Grossly Normal | + |  |
| A45 | *Pycnopodia helianthoides* | Dutch Harbor, Alaska | 2015-03-11 | Tube Foot | Grossly Normal | + |  |
| A46 | *Pycnopodia helianthoides* | Dutch Harbor, Alaska | 2015-03-11 | Tube Foot | Grossly Normal | + |  |
| A47 | *Pycnopodia helianthoides* | Dutch Harbor, Alaska | 2015-03-11 | Tube Foot | Grossly Normal | + |  |
| A48 | *Pycnopodia helianthoides* | Dutch Harbor, Alaska | 2015-03-11 | Tube Foot | Grossly Normal | + |  |
| A49 | *Pycnopodia helianthoides* | Dutch Harbor, Alaska | 2015-03-11 | Tube Foot | Grossly Normal | + |  |
| A50 | *Pycnopodia helianthoides* | Dutch Harbor, Alaska | 2015-03-11 | Tube Foot | Grossly Normal | + |  |
| A69 | *Pycnopodia helianthoides* | Dutch Harbor, Alaska | 2015-03-11 | Tube Foot | Grossly Normal |  |  |
| A72 | *Pycnopodia helianthoides* | Dutch Harbor, Alaska | 2015-03-11 | Tube Foot | Grossly Normal | + |  |
| A74 | *Pycnopodia helianthoides* | Dutch Harbor, Alaska | 2015-03-11 | Tube Foot | Grossly Normal |  |  |
| A80 | *Pycnopodia helianthoides* | Dutch Harbor, Alaska | 2015-03-11 | Tube Foot | Grossly Normal | + |  |
| A84 | *Pycnopodia helianthoides* | Dutch Harbor, Alaska | 2015-03-11 | Tube Foot | Grossly Normal |  |  |
| A85 | *Pycnopodia helianthoides* | Dutch Harbor, Alaska | 2015-03-11 | Tube Foot | Grossly Normal | + |  |
| A87 | *Pycnopodia helianthoides* | Dutch Harbor, Alaska | 2015-03-11 | Tube Foot | Grossly Normal | + |  |
| A96 | *Pycnopodia helianthoides* | Dutch Harbor, Alaska | 2015-03-11 | Tube Foot | Grossly Normal |  |  |
| C511 | *Pycnopodia helianthoides* | Puget Sount | 2016-01-10 | Coelomic Fluid | Grossly Normal | + |  |
| C555 | *Orthasterias kohleri* | Puget Sount | 2016-01-10 | Coelomic Fluid | Grossly Normal |  |  |
| C605 | *Dermasterias imbricata* | Puget Sount | 2016-01-11 | Coelomic Fluid | Grossly Normal |  |  |

PCR amplicons were run on 2% agarose gels in 1X TBE for 60 min at 100V, including the 100 bp ruler. Gels were stained in 1X SYBR Gold or SYBR Green for 30 min, and visualized in a BioRad ChemDoc system. Reactions bearing amplicons at the predicted 321 nt product for Vpec_F/Vpec_R and at 250 nt for V4 16S rRNA were scored as positives, while those for which no amplicon was visible under greatest transilluminator intensity were scored as negative. Four Vpec_F/Vpec_R amplicons were haphazardly selected for Sanger sequencing to confirm target specificity.

### 3. PCR amplification of V. pectenicida in coelomic fluid of asteroids that had SSW signs during experimental incubations

Samples of coelomic fluid collected during an experiment to examine microbiome changes during organic matter enrichment, performed at the Bodega Bay Laboratory in August 2019 (Aquino et al. 2021) and hitherto unreported were surveyed for the presence of *V. pectenicida* FHCF-3 (Table 4). Coelomic fluid samples, which were withdrawn from arms of each star every day during the course of the experiment, were selected to represent either specimens that never wasted over the course of the 15 d experiment (n = 4), or those which wasted after 4 to 12 days post treatment (n = 16). Specimens of coelomic fluid were targeted from the timepoint (24 h) immediately before wasting (body wall lesion) onset to distinguish potential etiologic agents from taxa colonizing decaying tissues. Coelomic fluid samples were thawed on ice, then 200 µl were extracted using the Zymo Tissue & Insect kit according to manufacturer’s recommendations. The extracted DNA was then subject to the V4 16S rRNA and Vpec_F/Vpec_R PCR amplifications as outlined above and visualized on electrophoretic gels.

**Table 4:** *Pisaster ochraceus* coelomic fluid samples consulted with Vpec_F/Vpec_R PCR for the presence of *Vibrio pectenicida* FHCF-3. Ampl V4 16S rRNA = Amplicon produced using V4 16S rRNA (prokaryotic-wide) primers; Ampl. Vpec_F/Vpec_R = Amplicon produced using the Vpec_F/Vpec_R primer pair. Note that PCR amplicons using Vpec_F/Vpec_R should be considered overestimates since amplicon sequences of several specimens were from *Vibrio* other than *V. pectenicida* FHCF-3.

| Asteroid No | Day | Date | Treatment | Tissue State | Ampl. V4 16S rRNA | Ampl. Vpec_F/Vpec_R |
| --- | --- | --- | --- | --- | --- | --- |
| 1 | 15 | 2019-08-15 | Control | Grossly Normal | + | + |
| 2 | 15 | 2019-08-15 | Control | Grossly Normal | + |  |
| 3 | 6 | 2019-08-06 | Control | Day Before Abnormal | + |  |
| 4 | 15 | 2019-08-15 | Control | Grossly Normal | + | + |
| 5 | 8 | 2019-08-08 | Control | Day Before Abnormal | + | + |
| 6 | 8 | 2019-08-08 | Peptone | Day Before Abnormal | + |  |
| 7 | 2 | 2019-08-02 | Peptone | Day Before Abnormal | + |  |
| 8 | 4 | 2019-08-04 | Peptone | Day Before Abnormal | + | + |
| 9 | 4 | 2019-08-04 | Peptone | Day Before Abnormal | + |  |
| 10 | 4 | 2019-08-04 | Peptone | Day Before Abnormal | + | + |
| 11 | 2 | 2019-08-02 | Dunaliella | Day Before Abnormal | + |  |
| 12 | 2 | 2019-08-02 | Dunaliella | Day Before Abnormal | + | + |
| 13 | 8 | 2019-08-08 | Dunaliella | Day Before Abnormal | + | + |
| 14 | 10 | 2019-08-10 | Dunaliella | Day Before Abnormal | + | + |
| 15 | 2 | 2019-08-02 | Dunaliella | Day Before Abnormal | + |  |
| 16 | 10 | 2019-08-10 | Coastal POM | Day Before Abnormal | + |  |
| 17 | 2 | 2019-08-02 | Coastal POM | Day Before Abnormal | + |  |
| 18 | 15 | 2019-08-15 | Coastal POM | Grossly Normal | + |  |
| 19 | 8 | 2019-08-08 | Coastal POM | Day Before Abnormal | + |  |
| 20 | 4 | 2019-08-04 | Coastal POM | Day Before Abnormal | + | + |

### 4. Consultation of 16S rRNA amplicon libraries from bacterioplankton and sediments

The *Vibrio pectenicida* FHCF-3 16S rRNA gene was compared by BLASTn against 16 bacterioplankton and 2 sediment bacterial amplicon libraries available in NCBI GenBank, representing estuarine and coastal sites (Table 5). Since several BioProjects included data from other compartments, only libraries meeting the criteria of bacterioplankton (or seawater) or sediments were included from these.

**Table 5:** Bacterioplankton and sediment bacterial libraries consulted against the *Vibrio pectenicida* FHCF-3 V4 16S rRNA amplicon sequence (copy 1 and 2) by BLASTn.

| Bioproject | Copy 1<br>matches<br>100% | Copy 2<br>matches<br>100% | Geographic Location | Target | Year | Citation |
| --- | --- | --- | --- | --- | --- | --- |
| PRJNA776096 | 28 | 92 | East Coast Australia | Bacterioplankton | 2020 | (Williams et al. 2022) |
| PRJNA1024631 | 47 | 37 | Moreton Bay - Brisbane River Estuary | Bacterioplankton | 2021 | n/a |
| PRJNA626452 | 13 | 13 | Water around New Zealand Shellfish Aquaculture | Bacterioplankton | 2015 | n/a |
| PRJEB14040 | 0 | 0 | Gulf of Naples, Italy | Bacterioplankton | 2013 | (Richa et al. 2017) |
| PRJNA514222 | 0 | 0 | Toulon Bay, France | Bacterioplankton | 2015 | (Coclet et al. 2019) |
| PRJNA1184006 | 0 | 0 | Changjiang River Estuary, China | Bacterioplankton | 2021 | n/a |
| PRJNA973211 | 0 | 0 | Pearl River Estuary | Bacterioplankton | 2017 | n/a |
| PRJNA951088 | 0 | 0 | Shrimp Aquaculture, Ningbo, China | Bacterioplankton | 2021 | (Zheng et al. 2023) |
| PRJNA927312 | 0 | 0 | Broadkill River, Lewes DE | Bacterioplankton | 2021 | (Guider et al. 2024) |
| PRJNA842171 | 0 | 0 | St Lawrence Waterway | Bacterioplankton | 2018 | (Jia et al. 2022) |
| PRJNA792736 | 0 | 0 | Paerl River Estuary | Bacterioplankton | 2017 | (Lu et al. 2023) |
| PRJNA775984 | 0 | 0 | Caloosahatchee River and Estuary | Bacterioplankton | 2022 | n/a |
| PRJNA684972 | 0 | 0 | Arno River Estuary | Bacterioplankton | 2015 | (Navarro et al. 2023) |
| PRJNA670217 | 0 | 0 | Patagonian fjord | Bacterioplankton | 2017 | (Tamayo-Leiva et al. 2021) |
| PRJNA593265 | 0 | 0 | Southern California Lagoons | Bacterioplankton | 2015 | (Diner et al. 2019) |
| PRJNA1233806 | 0 | 0 | Hong Kong Bay | Bacterioplankton | 2021 | (Wan et al. 2025) |
| PRJNA774107 | 0 | 0 | Yangtze River Sediments | Sediments | 2020 | n/a |
| PRJNA734698 | 0 | 0 | Xijiang River outlet, China | Sediments | 2020 | n/a |

### 5. Response of Vibrio spp. to echinoderm tissue decay

To assess how microorganisms in plankton and within tissues respond to the decay of *Pisaster ochraceus* tissues, a mesocosm experiment was performed. Body wall tissues from a single frozen *P. ochraceus* specimen (SC-06, Santa Cruz, July 2018; (Aquino et al. 2021)) was thawed and divided into 12 pieces (5 by 5 mm^2^, 0.05 g each). Half of the pieces were autoclaved for 1 h to kill remaining organisms present after freezing at −20°C for ∼8 years, while six specimens remained unsterilized. Ten liters of seawater from the Woods Hole Oceanographic Institution aquarium room intake were filtered through 0.2 µm Durapore filters to remove bacteria. Mesocosms (1L) were established containing: filtered water (n = 3); unfiltered water (n = 3); filtered water and sterilized tissue (n = 3); unfiltered water and sterilized tissue (n = 3); filtered water and unsterilized tissue (n = 3); and unfiltered water and unsterilized tissue (n = 3). Mesocosms were mixed once daily using a sterile serological pipette. Plankton samples were collected daily by filtering 50 – 200 mL water samples through 0.2 µm Durapore membranes using a syringe filter. The tissue piece was also removed using clean forceps and placed on a sterile petri dish where it was sampled (approx. 2 mm^2^) using 4 mm sterile biopsy punches. DNA was extracted from tissue pieces and 0.2 µm Durapore filters using the Zymo Tissue and Insect kit following manufacturer’s recommendations. DNA was quantified by Pico Green fluorescence, and diluted to 0.1 ng uL^-1^. Diluted DNA was submitted to the Michigan State University Genomics Core for V4 16S rRNA amplicon library preparation and sequencing. All specimens were sequenced on a single run of an Illumina MiSeq instrument. Sequence data is available at NCBI under accession PRJNA1306195.

Amplicon sequences were first imported into Qiita (which is based on the Qiime platform) (Caporaso et al. 2010), trimmed for length (250 nt) and subject to Deblur (Reference phylogeny for SEPP: Greengenes_13.8). From there, reads were subject to a pre-fitted sklearn-based taxonomy. The resulting biom file was parsed for “*Vibrio*” and resulting annotations imported as a csv file into Microsoft Excel. Matches to “*Vibrio*” were summed and expressed as a proportion of total reads in samples.

## Results

### 1. Survey of echinoderm samples and sequence databases from earlier mass mortality event

Comparison of the *Vibrio pectenicida* FHCF-3 16S rRNA sequence copy 1 (0178.1) against 41 RNA viromes prepared from asteroids and holothurians collected in 2013 – 2015 yielded 569 idential reads, with most hits in the southwest Pacific holothurian *Holothuria scabra* (n = 43; Moreton Bay, Australia, 2015), abnormal *Pisaster ochraceus* from Santa Cruz, CA (n = 498; abnormal in the absence of external stimuli 2018), and abnormal *Pyncopodia helianthoides* from the Seattle Aquarium (n = 12; 2013) (Table 1). Transcriptomes prepared from *P. helianthoides* yielded a larger number of reads matching *V. pectenicida* FHCF-3 copy 1 than viromes, with greatest recovery from body wall tissues prepared from grossly normal and abnormal specimens from the Seattle Aquarium in 2014 (>5,000 reads) (Gudenkauf & Hewson, 2015) and from coelomocytes of specimens treated with 0.2 µm filtered tissue homogenates of abnormal tissues (n = 995 reads) (Fuess et al. 2015). Assembly of read matches within each library resulted in 30 assembled contigs >200 nt. Comparison of these contigs against the core_nt library at NCBI and aligned contigs against close *Vibrio* relatives retrieved from NCBI discarded 18 that could not be unambiguously assigned to *V. pectenicida* FHCF-3 (i.e. they matched most closely to *Vibrio* spp. or unrelated bacteria (n = 10) or were ambiguous matches to conserved regions (n= 8)). The remaining contigs (n = 10; Table 6) matched at 100% nucleotide identity to *V. pectenicida* FHCF-3 16S rRNA gene copies 1, 4, 6 or 7. These contigs were from grossly normal and abnormal *Pycnopodia helianthoides* body wall and other tissue transcriptomes (n = 2), abnormal *P. helianthoides* coelomic fluid transcriptomes (n = 1), abnormal ray cross section *Pycnopodia helianthoides* RNA and DNA viromes (n = 2), abnormal *Pisaster ochraceus* ray cross section RNA and DNA viromes (n=2), a *Holothuria scabra* body wall RNA virome (n = 1), an *Astropecten polyacanthus* ray cross section DNA virome (n = 1), and a DNA virome of a ray cross section of *Echinaster luzonicus* (n = 1). Phylogenetic analyses of overlapping sequences placed these contigs into two groups: the first with abnormal *Pisaster ochraceus* body wall (in the absence of external stimuli) RNA Viromes from Santa Cruz 2018 (Aquino et al. 2021), abnormal and grossly normal *Pycnopodia helianthoides* body wall transcriptomes (2013) (Gudenkauf & Hewson, 2015), and from the *Pycnopodia helianthoides* coelomocyte transcriptomes (2014) (Fuess et al., 2014) (Fig. 1), and the second with rRNA contigs from abnormal *Pycnopodia helianthoides* and *Pisaster ochraceus* from DNA viromes in 2013 (Hewson et al. 2014a), grossly normal *Leptasterias* sp. contigs from RNA viromes in 2017 (Jackson et al. 2022), *Pycnopodia helianthoides* transcriptomes prepared in 2020 (Schiebelhut et al. 2024), and both grossly normal and abnormal *Apostichopus californicus* collected from Ketchikan, AK in 2016 (Fig. 2).

**Fig. 1:**
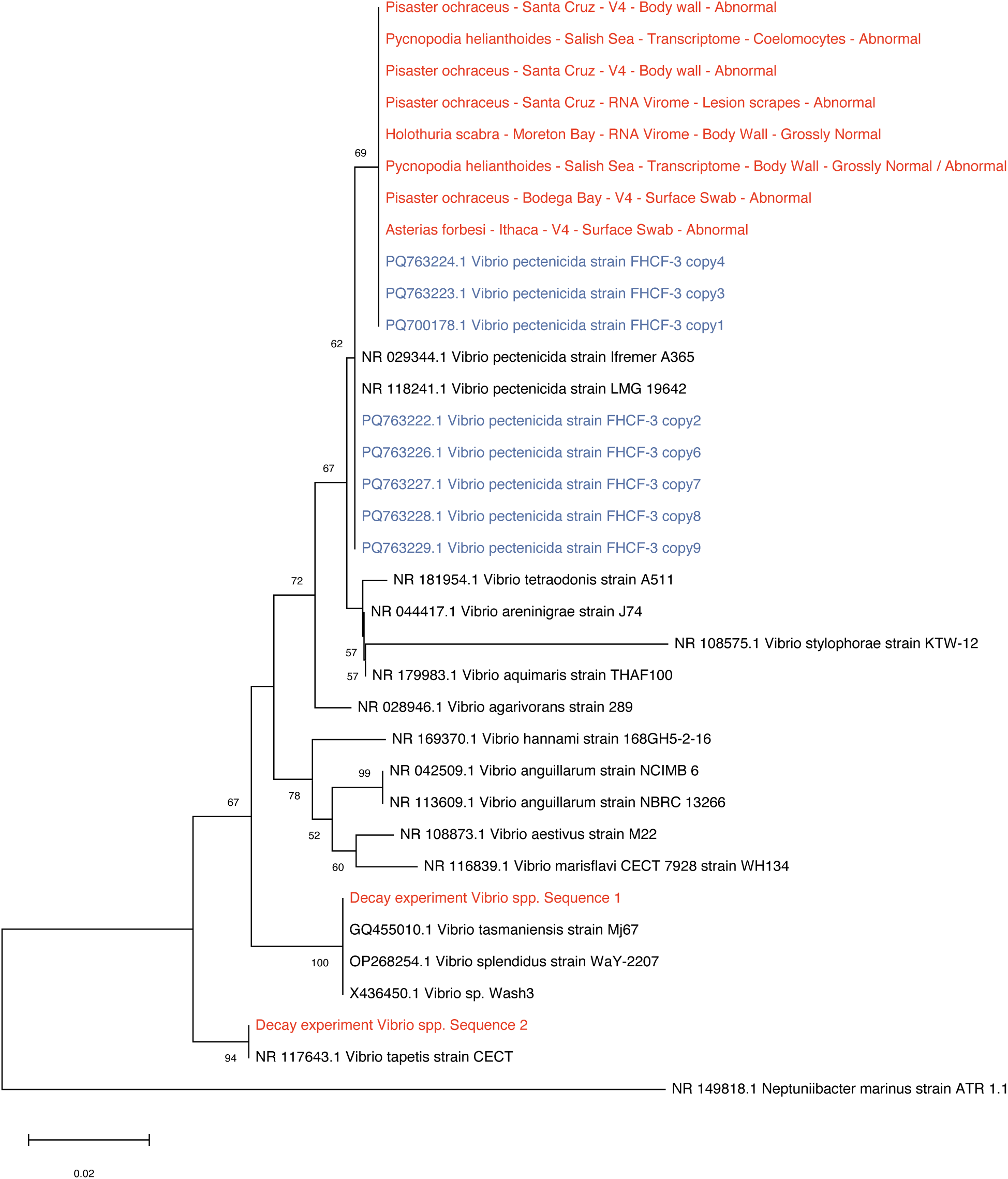
Phylogenetic reconstruction of the V4 region of 16S rRNA gene sequences derived in this study matching that region amongst contiguous sequences and 16S rRNA gene amplicon studies. Query sequences within variable regions were aligned against close relatives using MUSCLE (Edgar 2004), trimmed for non-overlapping alignment manually, and then subject to phylogenetic analysis using MEGAX (Kumar et al. 2018). Phylogenetic reconstructions were performed using the Jukes-Cantor and Neighbor Joining, with uniform rates of nucleotide replacement and 1000 bootstrap iterations. Red sequences are those recovered in this study; blue sequences are the 9 copies of *Vibrio pectenicida* FHCF-3.

**Fig. 2:**
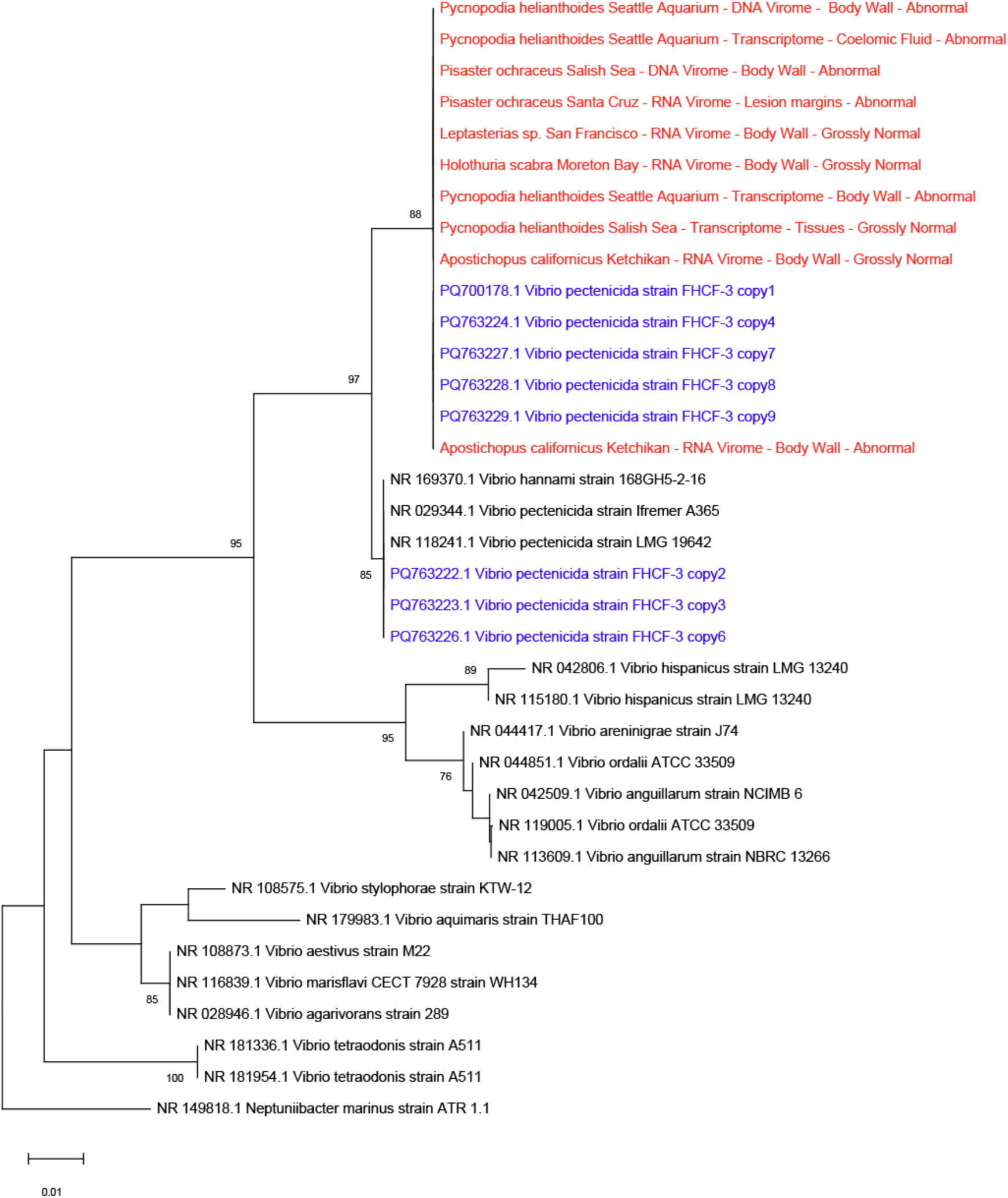
Phylogenetic reconstruction of 16S rRNA sequences derived in this study matching a region outside of the V4 16S rRNA amplicon. Query sequences within variable regions were aligned against close relatives using MUSCLE (Edgar 2004), trimmed for non-overlapping alignment manually, and then subject to phylogenetic analysis using MEGAX (Kumar et al. 2018). Phylogenetic reconstructions were performed using the Jukes-Cantor and Neighbor Joining, with uniform rates of nucleotide replacement and 1000 bootstrap iterations. Red sequences are those recovered in this study; blue sequences are the 9 copies of *Vibrio pectenicida* FHCF-3.

**Table 6:** Summary of BLASTn results against the NCBI core_nt database of contiguous 16S rRNA sequences >200nt reconstructed from 100% identical matches to *Vibrio pectenicida* FHCF-3 copy 1.

| Species | Length (nt) | Best match to core_nt | NCBI Accession No | % nucleotide identity | E-value |
| --- | --- | --- | --- | --- | --- |
| *Astropecten polyacanthus* | 211 | Vibrio pectenica strain FHCF-3 copy7 16S ribosomal RNA gene | PQ763227.1 | 100 | 4.00E-104 |
| *Holothuria scabra* | 1256 | Vibrio pectenica strain FHCF-3 copy1 16S ribosomal RNA gene | PQ700178.1 | 100 | 0 |
| *Pycnopodia helianthoides* | 279 | Vibrio pectenica strain FHCF-3 copy7 16S ribosomal RNA gene | PQ763227.1 | 100 | 8.00E-142 |
| *Echinaster luzonicus* | 378 | Vibrio pectenica strain FHCF-3 copy7 16S ribosomal RNA gene | PQ763227.1 | 100 | 0.00E+00 |
| *Pycnopodia helianthoides* | 1387 | Vibrio pectenica strain FHCF-3 copy1 16S ribosomal RNA gene | PQ700178.1 | 100 | 0 |
| *Pisaster ochraceus* | 251 | Vibrio pectenica strain FHCF-3 copy4 16S ribosomal RNA gene | PQ763224.1 | 100 | 3.00E-126 |
| *Pisaster ochraceus* | 1258 | Vibrio pectenica strain FHCF-3 copy1 16S ribosomal RNA gene | PQ700178.1 | 100 | 0 |
| *Pycnopodia helianthoides* | 550 | Vibrio pectenica strain FHCF-3 copy7 16S ribosomal RNA gene | PQ763227.1 | 100 | 0 |
| *Tigriopus californicus* | 290 | Vibrio pectenica strain FHCF-3 copy6 16S ribosomal RNA gene | PQ763226.1 | 100 | 6.00E-148 |
| *Pycnopodia helianthoides* | 1522 | Vibrio pectenica strain FHCF-3 copy1 16S ribosomal RNA gene | PQ700178.1 | 100 | 0 |

Comparison of the *V. pectenicida* 16S rRNA sequence copy 1 against V4 16S rRNA amplicons yielded hits to libraries prepared from surface swabs of *Pisaster ochraceus* tissues treated with organic matter substrates (10,592 reads), and body wall samples of *Pisaster ochraceus* that developed lesions over time in the absence of external stimuli in aquaria and which were sampled several days after lesions appeared (2,543 reads) (Table 2) (Aquino et al. 2021). A further library that sampled *Pisaster ochraceus* lesion margins and tissue scrapes away from lesions (in the absence of external stimuli) and were sampled at the time of lesion development yielded fewer reads (n=53) (Aquino et al. 2021). Reads matching *V. pectenicida* 16S rRNA copy 1 were not recovered from tissues of asteroids collected in January 2016 from the Salish Sea, from tissues of various asteroid species collected in Moreton Bay and Heron Island, Australia in December 2015 (Jackson et al. 2018), or from tissues of *Apostichopus californicus* during an organic matter amendment experiment in November 2021 (Crandell et al. 2023).

Comparison of *V. pectenicida* 16S rRNA copy 1 against the transcriptome shotgun assembly (TSA) database at NCBI yielded 2 reads matching a transcriptome of *Tigriopus californicus* prepared prior to 2012 (Schoville et al. 2012) from specimens collected from Santa Cruz CA and grown to high density (>300 specimens) in a culture facility prior to preparation.

### 2. PCR amplification of V. pectenicida in coelomic fluid of asteroids that had SSW signs during experimental incubations

PCR amplification of V4 16S rRNA gene from 91 specimens of various abnormal and normal asteroids collected in 2013 – 2015 yielded 72 samples that had viable DNA (the remainder were likely degraded due to the advanced age of extracts) (Table 3). Of these, 21 amplified by PCR employing the Vpec_F/Vpec_R primer set designed in this study. These were primarily from abnormal *Pycnopodia helianthoides* (n = 18 of 21 viable extracts) collected from the Seattle and Vancouver aquariums in 2013-2014 (body wall and pyloric caeca samples). One abnormal specimen from wild *Pisaster ochraceus* body wall (Santa Cruz; n = 20 abnormal and 2 grossly normal), and two *Evasterias troscheli* (one body wall and pyloric caeca) from the Vancouver Aquarium (n = 6 abnormal; n = 1 grossly normal) also yielded Vpec_F/Vpec_R amplicons.

Because amplicon quantity for these latter three was poor, specimens were subjected to a second PCR amplification using original template material, but these failed to yield further amplicons. PCR amplification of Vpec_F/Vpec_R also was unsuccessful in 25 grossly normal *Pycnopodia helianthoides* tube foot specimens from a reference site in Dutch Harbor Alaska (March 2015; Table 3). Four amplicons from abnormal *Pycnopodia helianthoides* body wall (Seattle Aquarium, October 2013) were subject to Sanger sequencing. All four sequences were phylogenetically associated with other *Vibrio pectenicida* strains in the Mediterranean and *Vibrio* spp. sequences derived from animal tissues and an intertidal mud flat, distinct from *Vibrio pectenicida* strain FHCF-3 (all copies) (Fig. 3).

**Fig 3:**
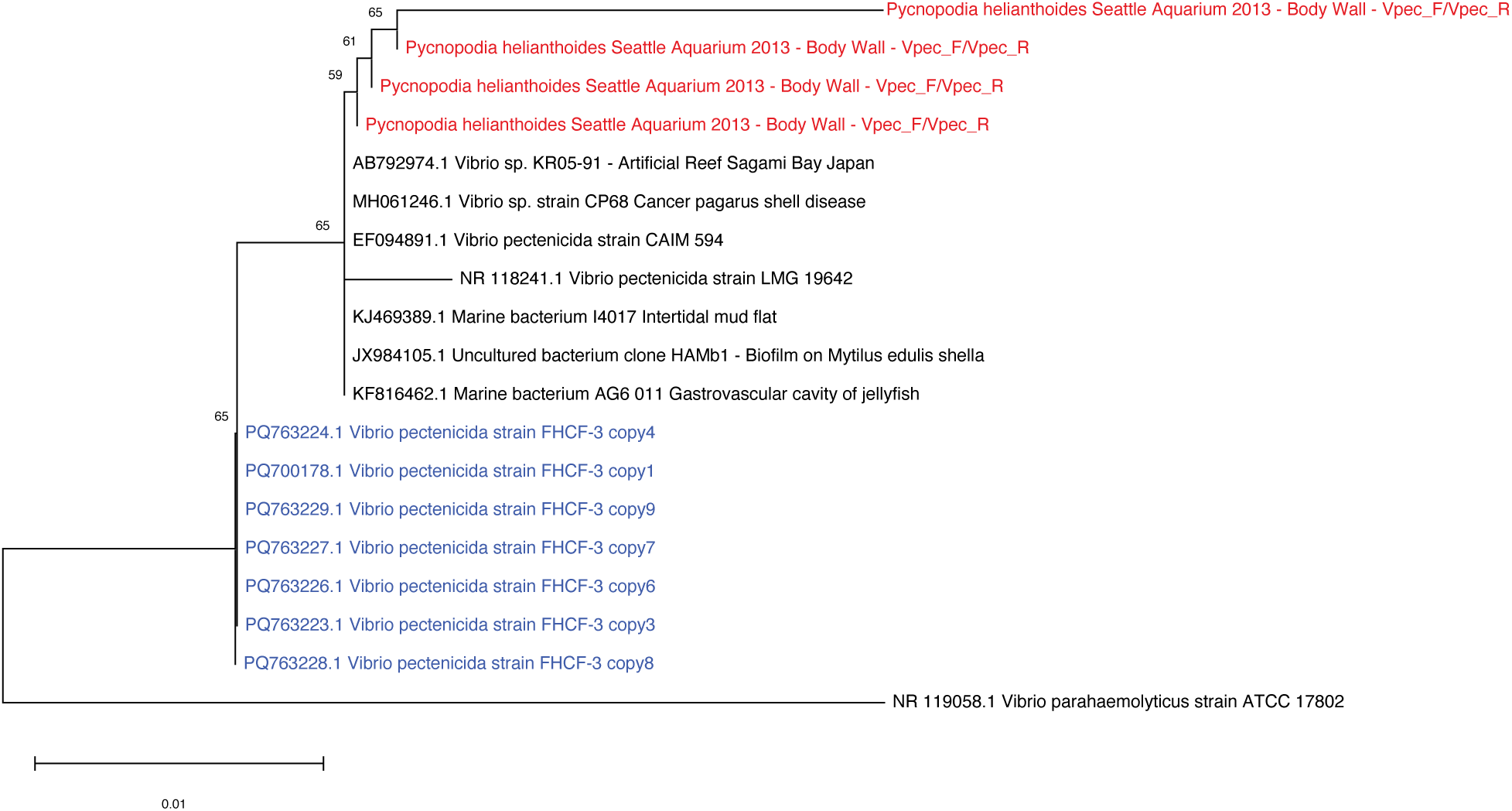
Phylogenetic relationship of PCR-amplified 16S rRNA fragments using primers Vpec_F/Vpec_R from abnormal Seattle Aquarium specimens collected in 2013. Query sequences within variable regions were aligned against close relatives using MUSCLE (Edgar 2004), trimmed for non-overlapping alignment manually, and then subject to phylogenetic analysis using MEGAX (Kumar et al. 2018). Phylogenetic reconstructions were performed using the Jukes-Cantor and Neighbor Joining, with uniform rates of nucleotide replacement and 1000 bootstrap iterations. Red sequences are those recovered in this study; blue sequences are the 9 copies of *Vibrio pectenicida* FHCF-3.

### 3. PCR amplification of V. pectenicida FHCF-3 16S rRNA in coelomic fluid of asteroids that had SSW signs during experimental incubations

In an experiment to test the impacts of organic matter loading on occurrence of SSW abnormalities (Aquino et al., 2021), 4 of 20 *Pisaster ochraceus* enrolled in the study remained grossly normal, while the remaining 16 developed body wall lesions 4 – 14 d after the experiment commenced. Coelomic fluid collected from the day prior to lesion observation from 1 *P. ochraceus* that never developed body wall lesions (a control) yielded PCR amplicons for VPec_F/Vpec_R, while 8 of 16 specimens that developed lesions yielded amplicons (Table 4).

To explore the relationship between SSW abnormalities and *V. pectenicida* FHCF-3 on animal surfaces, we recruited the 16S rRNA gene copy against each V4 16S rRNA gene library prepared from surface swabs separately. Abnormal specimens bore a greater percentage of *V. pectenicida* FHCF-3 copy 1 on their surface tissues the day before gross abnormal signs compared to those that did not become abnormal during the experiment (Fig. 4). The amendment with all 3 organic matter substrates (peptone, *Dunaliella* culture, and coastal POM) resulted in greater *V. pectenicida* FHCF-3 on abnormal specimen surfaces than control asteroids, with the greatest response to coastal DOM (Fig. 5). Comparison of coelomic fluid V_pecF/Vpec_R PCR detection with surface swab read proportion of 16S rRNA amplicon libraries revealed an inverse relationship, where coelomic fluid PCR positive samples bore fewer surface reads than those in PCR negative samples (Fig. 6).

**Fig. 4:**
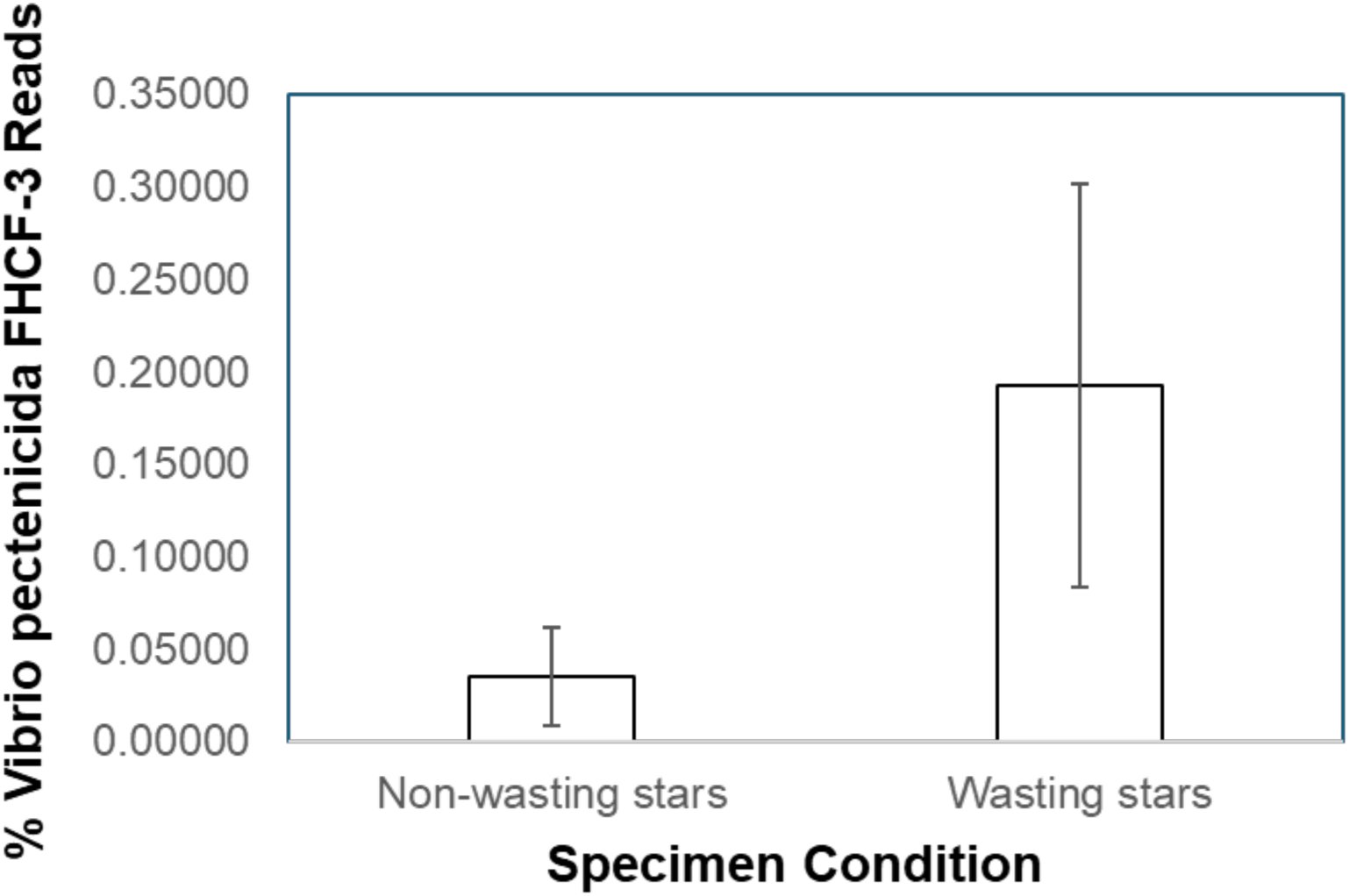
Comparison of *Vibrio pectenicida* FHCF-3 16S rRNA gene read recovery in surface swabs of asteroids that became abnormal during the experiment (24 h before appearance of lesions) and specimens that remained grossly normal throughout the experiment.

**Fig. 5:**
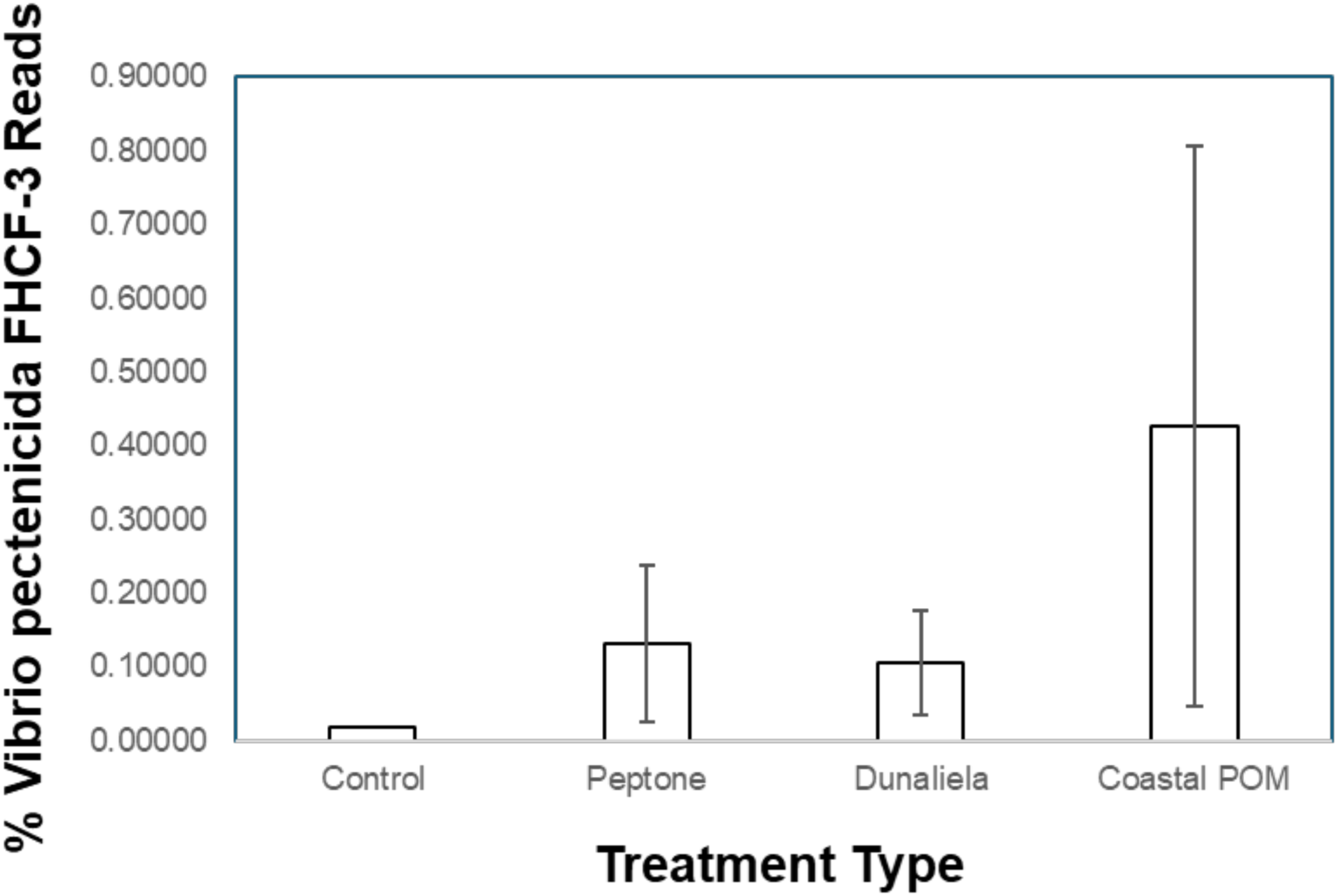
Effect of organic matter amendment on *Vibrio pectenicida* FHCF-3 read proportion in specimens that became abnormal during the experiment.

**Fig. 6:**
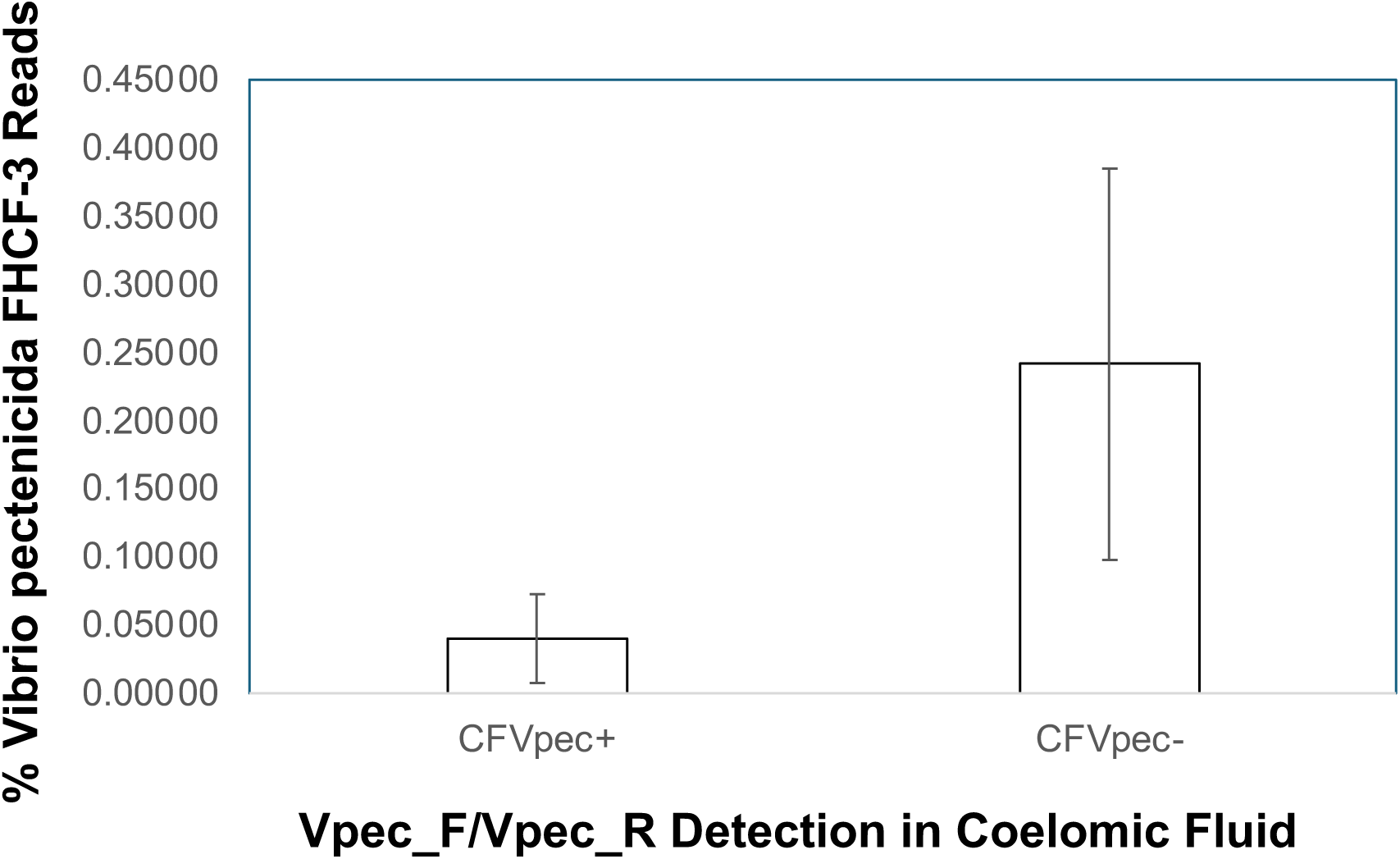
Comparison of Vpec_F/Vpec_R detection in coelomic fluid with proportion of *Vibrio pectenicida* FHCF-3 reads within amplicon libraries prepared from surface swabs. CFVpec+ = PCR amplification yielded amplicon using primers Vpec_F/Vpec_R; CFVpec-= PCR amplification did not yield amplicon using primers Vpec_F/Vpec_R.

### 4. Consultation of 16S rRNA amplicon libraries from bacterioplankton and sediments

The *Vibrio pectenicida* FHCF-3 16S rRNA gene copies 1 and 2 were retrieved from three surveys of free-living bacterioplankton from the southwestern Pacific Ocean (Table 5). These were to a survey of bacterioplankton along coastal sites of Eastern Australia (Williams et al. 2022), bacterioplankton in the Brisbane River and Moreton Bay in February 2021 (PRJNA1024631) and to plankton collected around shellfish aquaculture in Aotearoa New Zealand (PRJNA626452). *V. pectenicida* FHCF-3 was not detected in the remaining bacterioplankton and sediment bacteria surveys.

### 5. Response of Vibrio spp. to echinoderm tissue decay

The initial relative abundance of *Vibrionales* bacteria in mesocosms which were amended with dead *Pisaster ochraceus* tissues was 14.7 ± 0.2% of plankton reads and 0.4 ± 0.2 % of unsterilized tissue reads (Fig. 7). After 48 h, *Vibrionales* bacterial reads surged to 73-88% of total plankton and 61-81% of tissue reads, with the greatest increase in unsterilized pieces of *Pisaster ochraceus* tissues in filtered SW. The *Vibrionales* reads were affiliated primarily with two ribotypes, one matching *Vibrio splendidus*, and one matching *Vibrio tapetis* (Fig. 1).

**Fig. 7:**
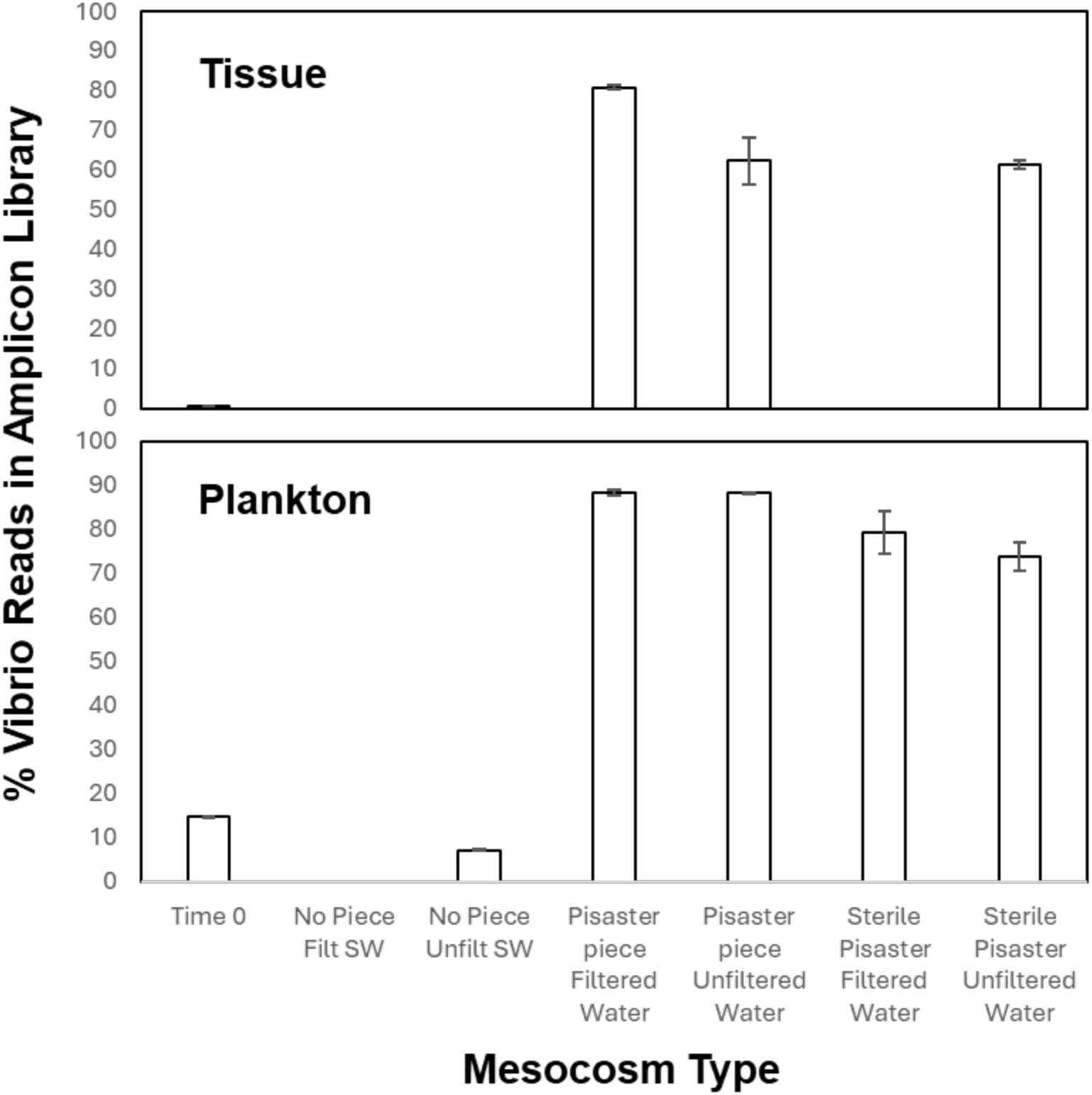
Impact of *Pisaster ochraceus* tissues on *Vibrio* spp. loads within tissues and plankton during a mesocosm study.

## Discussion

A key knowledge gap around *Vibrio pectenicida* FHCF-3 biology and ecology is its functional role in asteroid abnormalities consistent with gross signs of sea star wasting. This survey suggests that while *V. pectenicida* FHCF-3 was present in *Pycnopodia helianthoides* during both the initial asteroid mass mortality in fall 2013 and several experimental incubations of this species and *Pisaster ochraceus*, these data also do not support its association with other species that were abnormal at the time. Rather, the discovery of 16S rRNA genes identical to *V. pectenicida* FHCF-3 in geographically disparate samples and even other echinoderm orders suggests that it may be cosmopolitan and common on and in echinoderm tissues. In *Pisaster ochraceus*, *Vibrio pectenicida* FHCF-3 does not correspond with abnormalities consistent with SSW in coelomic fluid, but rather correlates with its abundance on animal surfaces. Enrichment of this taxon on abnormal *Pisaster ochraceus* surfaces in response to organic matter amendment suggest that it may thrive in high nutrient conditions and on decaying asteroid tissues. This is consistent with observations from a mesocosm study which found that *Vibrio* spp, experiences explosive growth in the presence of dead and decaying *Pisaster ochraceus* tissues. Together, these data suggest that *V. pectenicida* FHCF-3 is primarily a saprobic microorganism. Re-examination of experiments performed in the early mass mortality that yielded SSW abnormalities with 0.2 µm tissue homogenates and resulted in the appearance of *V. pectenicida* FHCF-3 in treated specimens when cells were not added to coelomic fluid, suggests that it may be a commensalistic taxon that is highly responsive to organic matter inputs. These facets raise interesting questions about its role in gross abnormalities consist with sea star wasting.

### 1. Vibrio pectenicida FHCF-3 is not intimately associated with tissues or pyloric caeca of wasting specimens of species other than Pycnopodia helianthoides

We approached detection of *Vibrio pectenicida* FHCF-3 in specimens from the 2013 mass mortality through two means. First, we interrogated DNA and RNA virome libraries prepared from tissues of affected and unaffected asteroids. Processing of homogenized tissues includes filtration through 0.2 µm filters; these are then treated with nucleases to reduce nucleic acids not within capsids or subcellular components like ribosomes (Thurber et al. 2009, Ng et al. 2011).

Despite these libraries targeting the virus-sized material, viral metagenomic libraries are frequently dominated by bacteria and eukaryotes and offer a potential source of detection for candidate pathogens (Hewson et al. 2024a). The DNA viral metagenomes prepared in 2014 (Hewson et al. 2014) were constructed from ray cross-sections, including body wall, gonad, pyloric caeca and coelomic fluid. The utility of surveying 16S rRNA and 18S rRNA genes in metagenomes of this size fraction to describe microbial autecology has been demonstrated previously (Hewson & Sewell 2021, Hewson et al. 2024a). Second, we interrogated transcriptomes prepared from both coelomocytes and body wall as reported previously (Gudenkauf & Hewson, 2015, Fuess et al. 2015, Schiebelhut et al. 2024). Finally, we applied a newly-developed PCR primer set (Vpec_F and Vpec_R) to broadly survey DNA extracts from body wall, pyloric caeca and coelomic fluid samples of both wild and captive asteroid specimens collected in 2013-2015.

PCR amplification using the Vpec_F/Vpec_R primer set generated amplicons across a number of specimens from 2013-2014. However, Sanger sequencing of a small number of these amplicons yielded sequences that were not identical to *Vibrio pectenicida* FHCF-3 16S rRNA copies, with closest matches in the core_nt database to other strains of *Vibrio pectenicida* and *Vibrio* spp. sequences recovered from animal surfaces and an intertidal mud flat. Hence, PCR amplification resulting in amplicons reported in Table 3 may include false positives (representing detections of related *Vibrio* spp.) and therefore should be interpreted conservatively. Furthermore, these data suggest that the lack of Vpec_F/Vpec_R PCR amplification, while matching at 100% nucleotide identity to the FHCF-3 strain in the priming region, are likely true negative detections.

Our results demonstrate that *V. pectenicida* FHFC-3 16S rRNA genes could be retrieved from specimens of *Pycnopodia helianthoides* (body wall, pyloric caecum, and coelomic fluid samples) in 2013-2014, as well as in *Pisaster ochraceus* body wall samples from Santa Cruz, CA in September 2013. However, Vpec_F/Vpec_R PCR amplification and virome recruitment was unsuccessful in other species, and notably absent from specimens from the earliest mass mortality observed along the Pacific coast of the Olympic Peninsula (Hewson et al. 2014a). Sea star wasting is described as affecting over 20 species of asteroids across the Northeastern Pacific Ocean. Since only *Pycnopodia helianthoides* was tested in the Prentice et al.(2025) study, and there is evidence of inconsistent effects on other species (Crandall et al. 2024), these results suggest that *Vibrio pectenicida* FHCF-3 is not tightly associated with abnormalities in other species. One possibility is that there was insufficient coelomic fluid in the ray cross-section samples to account for its detection. However, the *V. pectenicida* FHCF-3 16S rRNA gene was also detected in many body wall specimens from the time, suggesting that this approach is viable for detection.

*V. pectenicida* FHCF-3 was also detected in several aquarium-based 16S rRNA gene amplicon surveys in both tissue samples and external swabs (Aquino et al. 2021). Notably, the V4 sequence of *Vibrio pectenicida* FHCF-3 was recovered from several studies performed to assess the impact of organic matter on *Pisaster ochraceus* abnormalities (Bodega Marine Laboratory, 2019), abnormalities of *Pisaster ochraceus* during wasting in the absence of external stimuli (Long Marine Lab, 2018), and *Asterias forbesi* response to depleted oxygen (Cornell University, 2019). The enrichment of this taxon in response to multiple stimuli and during longitudinal study of abnormalities demonstrates its flexibility in aquarium studies. *Vibrio pectenicida* is a facultative anaerobic taxon (Lambert et al. 1998) which requires organic matter to proliferate, a scenario achieved during the addition of peptone, *Dunaliella salinicola*, and coastal POM in Bodega Marine Laboratory in 2019. Previous work has cited the role of microorganisms at the animal-water interface as driving sea star wasting (Aquino et al. 2021), and bacterial abundances on sea stars immediately prior to wasting are significantly higher than on grossly normal sea stars. Strictly anaerobic bacteria also appear prior to wasting onset, suggesting that the surface environment of asteroids is hypoxic (Aquino et al. 2021). These results confirm that *Vibrio pectenicida* FHCF-3 is one of a number of surface-associated bacteria that thrives on asteroid surfaces preceding the development of gross signs (Lloyd & Pespeni 2018, McCracken et al. 2023, Prentice et al. 2025), and proliferates when gross lesions manifest, possibly due to decaying tissues.

To distinguish between the roles of surface-associated and coelomic-fluid borne *V. pectenicida* FHCF-3 in sea star wasting, we also surveyed its presence in coelomic fluid specimens of all 20 *Pisaster ochraceus* from an experiment testing the impacts of organic matter loading on gross abnormalities consistent with sea star wasting (Aquino et al. 2021). We chose specimens of coelomic fluid from the day before the appearance of lesions to distinguish potential causative agents from saprobic organisms that may infiltrate compromised tissues through body wall erosions. Given that affected sea stars in the Prentice et al. (2025) study lost limbs 4 – 6 days after inoculation with cultured *Vibrio pectenicida* FHCF-3, it is reasonable to expect detectability in their fluids 24 h prior to the onset of wasting. Vpec_F/Vpec_R PCR amplification on DNA extracted from coelomic fluid of specimens that remained grossly normal throughout the experiment yielded 1 positive specimen, while only 7 of the 16 specimens that became abnormal over the course of the experiment yielded Vpec_F/Vpec_R amplicons (9 abnormal specimens had no detectable *V. pectenicida* FHCF-3 via this approach). These results suggest that in *Pisaster ochraceus*, *V. pectenicida* FHCF-3 in coelomic fluid may not be intimately tied to sea star wasting disease, and its presence on most specimen surfaces over time suggests that it may recruit to echinoderms in aquarium systems.

These results also suggest that organic matter amendment to the surfaces of *Pisaster ochraceus* may influence the relative abundance of *V. pectenicida* FHCF-3, since all amendments resulted in a greater proportion of reads in abnormal asteroids compared to those that received no amendment. The inverse relationship between read proportion in surface swabs and detection of *V. pectenicida* FHCF-3 in coelomic fluid may indicate that abnormalities observed on body wall tissues may be unrelated to the basal-to-surface processes observed in Work et al. (2021). Since coelomic fluid *V. pectenicida* FHCF-3 detection was inconsistently related to abnormalities, this may indicate that organic matter load, including that released from echinoderms and from exogenous sources, may foster its growth on surfaces. Furthermore, the growth of *V. pectenicida* FHCF-3 may contribute to the hypothesized oxygen depletion at this interface or via bioactive molecule production (e.g. enzymes) (Aquino et al. 2021) that may cause inflammation at its surface.

### 2. V. pectenicida was also observed in sea stars in east Asia, copepod culture, and bacterioplankton in Australia and Aotearoa New Zealand

*Vibrio pectenicida* FHCF-3 16S rRNA was recovered from several holothurian and other asteroid species. For example, recovery of this taxon’s 16S rRNA gene in both grossly normal and abnormal *Apostichopus californicus* (Holothuroida) specimens from the Ketchikan sea cucumber fishery in October 2016 concurs with its occurrence on sea stars at the time (Lloyd & Pespeni 2018). The *V. pectenicida* FHCF-3 16S rRNA gene was also recovered from two asteroid specimens from Hong Kong (*Astropecten polyacanthus*) and Okinawa (*Echinaster luzonicus*) in 2014 (Hewson et al. 2018), suggesting it may have a wider occurrence than the North American Pacific coast in 2013. This is also confirmed by recovery of *V. pectenicida* FHCF-3 from a holothurian from Moreton Bay, Australia (*Holothuria scabra*) (Hewson et al. 2020b) and from culture of the intertidal copepod *Tigriopus californicus* in 2010 (Schoville et al. 2012), preceding mass mortality by several years. Further detection of this taxon in plankton of eastern Australia and near shellfish mariculture in Aotearoa New Zealand suggests that it may be widespread in productive coastal environments, including those directly related influenced by mariculture. These results suggest that *V. pectenicida* FHCF-3 may occur in high productivity environments (e.g. sediments and culture conditions), concurring with its prevalence in aquarium-based challenge and organic matter amendment studies.

### 3. Vibrio pectenicida FHCF3 surged in experiments where tissue filtrates led to wasting in coelomic fluid, but not in heat-treated filtrates

A contrast to Prentice et al. (2025) comes from studies performed in 2014 where asteroid abnormalities resulted from injection of 0.2 µm filtered tissue homogenates and compared to heat-treated homogenates in challenge of *Pycnopodia helianthoides* (Hewson et al. 2014, Fuess et al. 2015). *V. pectenicida* FHCF-3 was detected in the (Fuess et al. 2015) study of coelomocytes only in treated *Pycnopodia helianthoides*; read hits to control libraries, once assembled and compared against core_nt yielded no confident detection of this bacterium. This is perplexing because both challenges did not amend coelomic fluid with bacterial cells, but rather organic material and viruses (i.e. materials passing through a 0.2 µm filter) derived from tissues. Heat treated controls generally bear lower concentrations of protein and overall DOM than untreated filtered homogenates (Hewson et al. 2024b), and it is unclear whether heating affects lability of echinoderm-derived OM. These results suggest that *V. pectenicida* FHCF-3 was introduced in the tested *Pycnopodia helianthoides*, perhaps through injection trauma, and that it responded to organic matter inputs or other materials in the filtered tissue homogenates to generate wasting conditions. Injection controls with filtered seawater, which lack the organic material in tissue homogenates, may not yield the same results (Prentice et al. 2025). Hence, caution should be taken in interpreting results of inoculation with both tissue homogenates and cultured cells against filtered or heat-treated controls, since these may foster the growth of normally surface-bound microorganisms introduced into the coelomic cavity through trauma.

### 4. Vibrio spp. bacteria related to V. pectenicida surge in response to dead asteroid tissues

*Vibrio* spp. are invariably described as copiotrophic taxa because they are often present in plankton at low abundances but comprise large proportions of plankton under high productivity settings (e.g. during algal blooms) (Gilbert et al. 2012, Westrich et al. 2016). A key question in disease ecology is the distinguishment of saprobic microorganisms (i.e. those consuming organic matter released from dead or moribund tissues) and opportunistic or even obligate pathogenic agents. We examined the impacts of dead *Pisaster ochraceus* tissues on *Vibrio* spp. relative abundances in plankton and within tissues themselves in a controlled and replicated experiment using seawater obtained from the intake of the Woods Hole Oceanographic Institution aquarium facility. The experimental design allowed us to assess whether microbial taxa normally associated with tissues or those recruited from plankton grow on decaying *Pisaster ochraceus* tissues (i.e. comparing autoclave-sterilized vs unsterilized and filtered to unfiltered seawater).

Our results strongly suggest that the related *Vibrio splendidus* and *Vibrio tapetis*, initially present in unfiltered water, experienced explosive growth in both tissues and plankton in response to decay. These results illustrate the very strong potential of related *Vibrio* spp. strains to grow on organic matter, especially those derived from *Pisaster ochraceus* tissues. It is worthwhile noting that while this study did not recover the *V. pectenicida* FHCF-3 16S rRNA gene, the *V. splendidus* strain was phylogenetically closest to *Vibrio* sp. Wash3, which was recovered from a sponge (*Suberites domuncula*) (Saidin et al. 2017), and that the group to which this bacterium belongs includes several described pathogenic microorganisms of fish and invertebrates. It is also interesting to note that in 1 abnormal field collected specimen (SCF0122), reads of *V. tapetis* exceeded those from *V. pectenicida* FHCF-3 in the Prentice et al. (2025) study.

### 5. A proposed mechanism of association between Vibrio pectenicida FHCF-3 and SSW

Several synergistic lines of evidence point to a potential role of *Vibrio pectenicida* FHCF-3 in SSW gross abnormalities, notably that: it appears to be responsive to organic matter amendment (Aquino et al., 2021); it thrives in decomposing tissues (e.g. body wall lesions) (Aquino et al. 2021); it is present sporadically and inversely to surface abundances in coelomic fluid (Aquino et al. 2021); it is associated with body wall lesions when tissue organic substrates are amended to the coelomic cavity of *P. helianthoides* with presumably commensal *V. pectenicida* FHCF-3 (Hewson et al. 2014b, Fuess et al. 2015); and that as an isolated microorganism, it can generate limb autotomy in *Pycnopodia helianthoides* when injected into the perivisceral cavity of grossly normal sea stars (Prentice et al. 2025). This role may not be as an invasive infectious agent, but rather as a driver or indicator of SSW abnormalities across species on the Pacific coast of North America.

Asteroids are biologically unlike most other marine invertebrates since their coelomic fluid supports large abundances of prokaryotes (10^4^ – 10^5^ cells mL^-1^), roughly 1 – 2 orders of magnitude less than surrounding seawater (Jackson et al. 2018). The composition of these prokaryotes are variable, with some species of asteroid maintaining coelomic fluid communities distinct from surrounding seawater, while others being most similar to those around them suggesting that there can be some selection by antimicrobial compounds or coelomocytes on microbial composition (Nakagawa et al. 2017). Because of this facet of their biology, establishing pathogenicity of any microorganism during challenge experiments crucially demands examination of echinoderm tissue-level (body wall and/or coelomocyte) changes in response to insults, and comparison against those described in field-collected specimens experiencing the condition in the wild. Given that prior work has found that SSW lesions are associated with edema, inflammation, and a basal-to-surface possible starting with ossicle inflammation in the absence of bacterial infiltration within tissues (Work et al. 2021), future work must compare histological and cytological findings with prior work to provide definitive evidence for *V. pectenicida* FHCF-3 as a pathogenic agent. It remains possible that *V. pectenicida* FHCF-3 has similar impacts on asteroids as other fast-growing *Vibrio* spp or even other copiotrophic microbes. Given its prominence on grossly normal specimen surfaces and within other tissues, detections within coelomic fluids of wild SSW abnormal specimens does not link experimental results to its role in disease process in the wild since it may be an efficient saprobic taxon degrading echinoderm tissues.

Furthermore, injecting this taxon directly into perivisceral coelomic fluids obscures transmission pathways in nature that might otherwise prevent its entry; in turn, because of its potential efficient use of echinoderm derived organic matter it may grow rapidly in this compartment, especially if injection also ruptured internal organs like gonads or pyloric caeca. To gain entry to perivisceral fluids of *Pycnopodia helianthoides* under typical conditions, microorganisms must either enter the coelomic cavity through the outer epithelium or through the madreporite and stone canal. Asteroids species vary in the mechanism by which coelomic fluid (and associated microorganisms) is replenished and rate of flow through madreporites (Ferguson 1994). *Pisaster ochraceus* has high rates of flow through this entry mechanism because its epithelium is relatively impermeable, whereas madreporite influx in *Pycnopodia helianthoides* is slow, and is primarily replenished through diffusion of water molecules across its outer epidermis, especially via tube feet and radial canals of the ambulacral fluid (Ferguson 1994). Hence, in *Pycnopodia helianthoides* microbial effects on specimens would be most prominent on its outer epidermis than within coelomic fluids, and vice versa for *Pisaster ochraceus*. The inconsistent detection of *Vibrio pectenicida* FHCF-3 in coelomic fluid of abnormal *Pisaster ochraceus* in this study, yet consistent association in *Pyncopodia helianthoides* reported in Prentice et al. (2025) is incongruent with comparative coelomic fluid flow.

I propose an alternative explanation of work presented in Prentice et al (2025) on the functional role of *Vibrio pectenicida* FHCF-3 in SSW that fits with experimental and field observations, and analyses performed in this study. Under typical conditions, *V. pectenicida* FHCF-3 appears to be a commensalistic microorganism inhabiting asteroid surfaces and coelomic fluids at low abundance during cooler months when primary production (and its associated supply of DOM to surrounding waters) and temperature are lowest, but increase in abundance as temperatures rise and rates of primary production increase into spring and summer. At peak seasonal primary production, elevated DOM concentrations and higher temperatures may lead to a proliferation of this taxon on and around asteroid surfaces. This elevated heterotrophic activity may lead to outer epithelial damage by *V. pectenicida* FHCF-3 and other heterotrophic taxa via an as-yet unidentified mechanism (e.g. extracellular enzymes, other bioactive macromolecules or respiratory diffusion limitation at the animal-seawater boundary; Aquino et al. 2021). Influx of this taxon and DOM into coelomic fluids may occur when their abundances on animal surfaces or in surrounding habitats reach critically high levels, stimulating growth of commensalistic *V. pectenicida* already in coelomic fluid and introducing new cells; these in turn may grow rapidly and generate stress responses like limb autotomy. This effect may become exacerbated by the presence nearby of decaying asteroid tissues, leading to a ‘snowball’ effect of SSW when asteroid population densities result in localized patches of enriched DOM. Ultimately, SSW may result from a combination of seasonal patterns of DOM inputs and temperature, a fast-growing heterotrophic and copiotrophic taxon that grows well on decaying asteroid tissues and DOM from primary production, and as-yet undefined mechanisms of generating tissue damage, eventual infiltration into the coelomic cavity and explosive growth there in some asteroid species (e.g. *Pycnopodia helianthoides*), resulting in stress responses.

It is unlikely that *Vibrio pectenicida* FHCF-3 acts alone as a driver of SSW, or that it is involved in all cases of SSW through time or worldwide. There is also no evidence that pathology results from invasive infection within tissues or indeed that there is consistent pathology between field and experimental abnormal specimens reported in Prentice et al. (2025). Hence, great care must be taken to avoid extrapolating results of the Prentice et al. (2025) study to the wider sea star wasting phenomenon until pathogenesis of *V. pectenicida* FHCF-3 is established, consistent tissue-level changes in response to this candidate agent are found, and that its ability to generate consistent pathology across the wide suite of affected asteroid species is assessed.

## Acknowledgements

Dan Buckley, Thierry Work, John Wares, and an anonymous reviewer provided helpful comments on an early manuscript draft. Lesanna Lahner, Martin Haulena, Lesanna Lahner, Martin Haulena, Stephen Fradkin, Nathaniel Fletcher, Colleen Burge, Peter Raimondi, Mitchell Johnson, Kalia Bistolas, Elliot Jackson and Jason Button provided field and aquarium samples analyzed in this work. The author is grateful to Rob Lampe for supply of seawater from the Woods Hole Oceanographic Institution aquarium. This work was supported by an award from the US National Science Foundation OCE-2049225.

